# The impact of gut microbiota on depressive-like behaviors and adult hippocampal neurogenesis requires the endocannabinoid system

**DOI:** 10.1101/718288

**Authors:** Grégoire Chevalier, Eleni Siopi, Laure Guenin-Macé, Maud Pascal, Thomas Laval, Aline Rifflet, Ivo Gomperts Boneca, Caroline Demangel, François Leulier, Gabriel Lepousez, Gérard Eberl, Pierre-Marie Lledo

**Affiliations:** Microenvironment and Immunity Unit, Institut Pasteur, INSERM U1224, Paris, France; Perception and Memory Unit, Institut Pasteur, CNRS UMR3571, Paris, France; Immunobiology of Infection Unit, Institut Pasteur, INSERM U1221, Paris, France; Biology and Genetics of Bacterial Cell Wall Unit, Institut Pasteur, Paris, France; Institut de Génomique Fonctionnelle de Lyon, Université de Lyon, Ecole Normale Supérieure de Lyon, CNRS UMR 5242, Lyon, France

## Abstract

Depression is the leading cause of disability worldwide. Recent observations have revealed an association between mood disorders and alterations of the intestinal microbiota, but causality remains yet to be established. Here, using unpredictable chronic mild stress (UCMS) as a mouse model of depression, we show that the UCMS mice display phenotypic alterations — characterized by an altered gut microbiota composition, a reduced adult hippocampal neurogenesis and a depressive-like behaviors — which could be transferred from UCMS donors to naïve recipient mice by fecal microbiota transplantation. The cellular and behavioral alterations observed in recipient mice were accompanied by a decrease in the endocannabinoid (eCB) signaling due to lower peripheral levels of fatty acid precursors of eCB ligands. The adverse effects of UCMS-transferred microbiota on adult neurogenesis and behavior in naïve recipient mice were alleviated by selectively enhancing the central eCB tone or by adding arachidonic acid, a fatty acid precursor of eCB ligands, to the diet. In the gut of both UCMS donors and recipients, the microbiota composition was characterized by a relative decrease in *Lactobacilli* abundance, and complementation of the UCMS recipient microbiota with a strain of the *Lactobacilli* genus was sufficient to restore normal eCB brain levels, hippocampal neurogenesis and to alleviate depressive-like behaviors. Our findings provide a mechanistic scenario for how chronic stress, diet and gut microbiota dysbiosis generate a pathological feed-forward loop that contributes to despair behavior via the central eCB system.

## INTRODUCTION

Depression is the leading cause of disability worldwide, currently affecting more than 300 million people^1^. Despite the prevalence of depression and its considerable economic impact, its pathophysiology remains highly debated. Yet, a better understanding of the mechanisms leading to depression is a prerequisite for developing efficient therapeutic strategies. However, unraveling the pathophysiology of depression is challenging, as depressive syndromes are heterogeneous and their etiologies likely to be diverse. Experimental and genetic studies have yielded several mechanisms including maladaptive responses to stress with HPA axis dysregulation, inflammation, reduced neuroplasticity, circuit dysfunctions and perturbation in neuromodulatory systems such as monoaminergic and endocannabinoid (eCB) systems.

A number of studies converge to indicate hippocampal alterations as critical in the pathogenesis of depression. For instance, hippocampal volume loss is a hallmark of clinical depression^2–4^. Likewise, rodent studies have demonstrated that chronic stress-induced depression impair adult hippocampal neurogenesis^5–8^. Furthermore, impaired hippocampal neurogenesis results in depressive-like behaviors in rodent, in part because hippocampal neurogenesis buffers the over-reactivity of the hypothalamic-pituitary-adrenal (HPA) axis in response to stress^9–11^. In that line, antidepressants and alternative antidepressant interventions stimulate adult hippocampal neurogenesis, which in turn dampens stress responses and restores normal behavior^12–14^. Adult hippocampal neurogenesis is thus considered as an important causal factor and a key marker of depression, although a direct causal link is still missing in human depression^15, 16^.

Over the last decade, the impact of the symbiotic microbiota on numerous host functions has been increasingly recognized. The wide variety of intestinal microbes affects many processes including immunity^17^, metabolism^18^ and the central nervous system^19^. In depressed patients, alterations in the composition of the intestinal microbiota (named dysbiosis) have been characterized^20, 21^. Furthermore, numerous studies on animal models have shown that the microbiota modulates anxiety^22–24^ and onset of neurological diseases associated to circuit dysfunctions^25, 26^ by releasing bacterial metabolites that can directly or indirectly affect brain homeostasis^19, 27^. In that line of ideas, microbiota from depressed patients alter behavior when transferred to antibiotic-treated rats^28^ and murine gut microbiota dysbiosis is associated with several neurobiological features of depression, such as low-grade chronic inflammation^29^, abnormal activity of the HPA axis^30, 31^ and decreased adult neurogenesis^12, 32^. The notion that microbiota is a critical node in the gut-brain axis is also supported by the observation that colitis, which depends on the gut microbiota, shows significant comorbidities with depression^33^. Finally, probiotic intervention has been shown to influence emotional behavior in animal models of depression^34–36^ and improve mood in depressive patients^37–39^. However, the molecular mechanisms linking intestinal microbiota and mood disorders remain largely unknown, partly due to the lack of experimental models.

To explore a causative role of the gut microbiota in stress-induced depressive behaviors, we used unpredictable chronic mild stress (UCMS), a mouse model of depression, and fecal microbiota transfer (FMT) from stressed donors to naïve mice. We found that the microbiota transplantation transmits the depressive behavioral symptoms, and reduces adult neurogenesis of the recipient mice. Metabolomic analysis reveals that recipient mice developed an altered fatty acid metabolism characterized by deficits in lipid precursors for eCBs, which resulted in impaired activity of the eCB system in the brain. Increase of the eCB levels after pharmacological blocking of the eCB degrading enzymes, or complementation of the diet with arachidonic acid, a precursor of eCBs, is sufficient to normalize both the microbiota-induced depressive-like behaviors and hippocampal neurogenesis in recipient mice. Lastly, our study reveals that UCMS induced a gut microbiota dysbiosis characterized by a decrease in *Lactobacilli* abundance also observed in recipient mice. Complementation of recipient mice with a strain of the *Lactobacilli* genus is sufficient to restore both eCB brain levels and hippocampal neurogenesis, alleviating the microbiota-induced despair behavior.

## RESULTS

### Transplantation of microbiota from stressed mice to naïve recipients transfers depressive-like behaviors and reduces adult neurogenesis

To establish a depressive-like state in mice, we submitted C57BL/6J mice for 8 weeks to UCMS, a well-defined mouse model of stress-induced depression^40–42^ (Fig. 1A and **Supplementary Table S1**). Consistent with previous reports, UCMS mice developed depressive-like behaviors, as shown by increased feeding latency in the “novelty suppressed feeding test” as compared to control mice (Fig. 1B), even though feeding drive was not affected (**Supplementary Fig. 1A**). This behavior reflects both anxiety and anhedonia. However, UCMS mice did not develop increased anxiety, as determined by the “light/dark box test” (Fig. 1C). Furthermore, UCMS mice showed increased grooming latency (Fig. 1D) and decreased self-grooming behavior in the “splash test” (**Supplementary Fig.1B and C**), reflecting symptoms of depression such as apathetic behavior^41^. The depressive-like state seen in UCMS mice was further confirmed in two prototypical tests for assessing depressive-like behaviors, the “tail suspension test” and the “forced swim test” (also named behavioral despair tests). UCMS mice showed increased immobility time in these two tests compared to control mice (Fig. 1E and F). We also observed that UCMS mice gained significantly less weight over time than control mice, as previously reported^43^ (**Supplementary Fig. 1D**). Altogether, these different behavioral tests demonstrate that 8 weeks of UCMS induce depressive-like but not anxiety-like behaviors in C57BL/6 mice.

**Figure 1.**
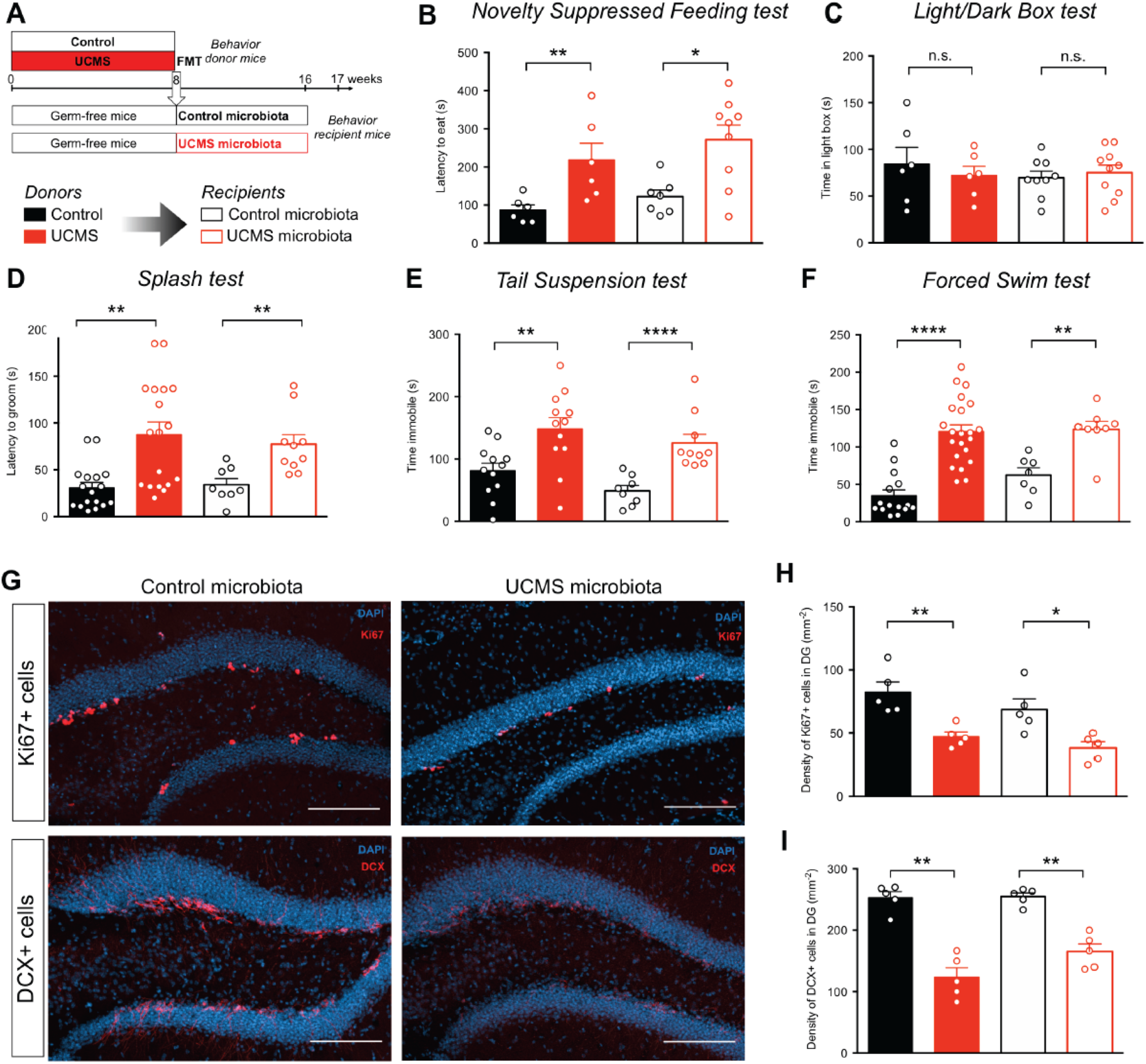
Microbiota from UCMS Mice Transfers Depressive-like Behaviors and Reduces Adult Hippocampal Neurogenesis. **A**, Experimental timeline of Fecal Microbiota Transplantation (FMT) from Control and UCMS mice, respectively ‘Control microbiota’ and ‘UCMS microbiota’, to germ-free recipient mice. **B-F,** Control mice (black bars), or mice subjected to UCMS (red bars), and mice recipient of the microbiota from Control (open black bars) or UCMS mice (open red bars), underwent different behavioral tests. **B,** Latency to eat in a novel environment in the Novelty Suppressed Feeding test for Control mice (*n* = 6), UCMS mice (*n* = 6), Control microbiota-recipient mice (*n* = 7) and UCMS microbiota-recipient mice (*n* = 9). (Control vs UCMS, *P* = 0.0087; Control microbiota- vs UCMS microbiota-recipient mice, *P* = 0.0229); **C**, Time spent in the light box in the Light/Dark Box test for Control mice (*n* = 6), UCMS mice (*n* = 6), Control microbiota-recipient mice (*n* = 9) and UCMS microbiota-recipient mice (*n* = 10). (Control vs UCMS, *P* = 0.6991; Control microbiota- vs UCMS microbiota-recipient mice, *P* = 0.6038); **D**, Latency to groom in the Splash test for Control mice (*n* = 17), UCMS mice (*n* = 18), Control microbiota-recipient mice (*n* = 8) and UCMS microbiota-recipient mice (*n* = 10). (Control vs UCMS, *P* = 0.0004; Control microbiota- vs UCMS microbiota-recipient mice, *P* = 0.0012); **E**, Time spent immobile in the Tail Suspension Test for Control mice (*n* = 10), UCMS mice (*n* = 10), Control microbiota-recipient mice (*n* = 8) and UCMS microbiota-recipient mice (*n* = 10). (Control vs UCMS, *P* = 0.0043; Control microbiota- vs UCMS microbiota-recipient mice, *P* < 0.0001); **F**, Time spent immobile in the Forced Swim Test for Control mice (*n* = 15), UCMS mice (*n* = 22), Control microbiota-recipient mice (*n* = 7) and UCMS microbiota-recipient mice (*n* = 8). (Control vs UCMS, *P* < 0.0001; Control microbiota- vs UCMS microbiota-recipient mice, *P* = 0.0037). **G**, Representative images of Ki67 staining (top) and DCX staining (bottom) in the DG of the hippocampus, counterstained with DAPI (blue), in Control microbiota-recipient mice (left) and UCMS microbiota-recipient mice (right). **H**, Quantitative evaluation of the density of Ki67^+^ cells in the dentate gyrus (DG) of the hippocampus for Control mice (*n* = 5), UCMS mice (*n* = 5), Control microbiota-recipient mice (*n* = 5) and UCMS microbiota-recipient mice (*n* = 5). (Control vs UCMS, *P* = 0.0079; Control microbiota- vs UCMS microbiota-recipient mice, *P* = 0.0159). Scale bar: 100µm. I, Quantitative evaluation of the density of DCX^+^ cells in the DG of the hippocampus for Control mice (*n* = 5), UCMS mice (*n* = 5), Control microbiota-recipient mice (*n* = 5) and UCMS microbiota-recipient mice (*n* = 5). (Control vs UCMS, *P* = 0.0079; Control microbiota- vs UCMS microbiota-recipient mice, *P* = 0.0079) For (**B** to **I**) Data are represented as mean ± s.e.m. Statistical significance was calculated using the Mann Whitney test (\**P* < 0.05, ** *P* < 0.01, *** *P* < 0.001, **** *P* < 0.0001).

As the reduction of adult hippocampal neurogenesis is a hallmark of depression, we tested whether UCMS affected the number of adult-born neurons in the dentate gyrus (DG) of the hippocampus. The decreased number of proliferating neural stem cells labeled with the cell proliferation marker Ki67 (Fig. 1G and H), and of doublecortin (DCX)^+^ cells, a marker for newborn immature neurons (Fig. 1G and I), shows that UCMS mice exhibit reduced hippocampal neurogenesis.

We next assessed whether the transplantation of gut microbiota from UCMS mice to naïve unstressed hosts was sufficient to transfer the hallmarks of depressive-like state. To this end, we transferred the fecal microbiota of control or stressed mice to adult germ-free mice (Fig. 1A). Eight weeks after FMT, recipients of UCMS microbiota showed depressive-like behaviors in both the tail suspension and the forced swim tests (Fig. 1E and F), which were confirmed in the splash test (Fig. 1D and **Supplementary Fig. 1F** and **G**) and the novelty suppressed feeding test (Fig. 1B and **Supplementary Fig. 1E**). As in UCMS donors, recipient mice did not express anxiety-related behaviors (Fig. 1C). Similar results were obtained when UCMS microbiota was transferred to recipient SPF mice that were treated with broad-spectrum antibiotics for 6 days until one day prior to FMT (**Supplementary Fig. 2**). Because germ-free mice might exhibit some behavioral abnormalities due to sustained disruption in the microbiota-gut-brain axis, all subsequent experiments were performed using short-term antibiotic-treated recipient mice. Finally, recipients of UCMS microbiota also showed decreased proliferation of neural stem cells (Fig. 1G and H) and decreased production of new neurons in the hippocampus (Fig. 1G and I). These data demonstrate that the hallmarks of depressive-like behaviors are transferable to naïve recipient mice by the transplantation of fecal microbiota obtained from stressed-induced depressive mice.

### Gut microbiota from stressed mice alters fatty acid metabolism and the hippocampal endocannabinoid system

We explored the possibility that UCMS microbiota triggered depressive-like behaviors through alterations of the host’s metabolism. Metabolomic profiling of serum revealed a significant decrease in the levels of monoacylglycerols (MAG) and diacylglycerols (DAG) in both UCMS mice and recipients of UCMS microbiota, as compared to control and recipients of control microbiota (Fig. 2A). Furthermore, the n-6 polyunsaturated fatty acid (PUFA), arachidonic acid (AA, 20:4n-6), its precursor linoleic acid (18:2n-6), and n6-PUFA biosynthesis intermediates, were significantly decreased in both UCMS donors and recipients (Fig. 2B and C). This lipids loss was specific to short-chain fatty acids since levels of several medium- and long-chain fatty acylcarnitines rather increased in UCMS microbiota recipients (**Supplementary Fig. 3**).

**Figure 2.**
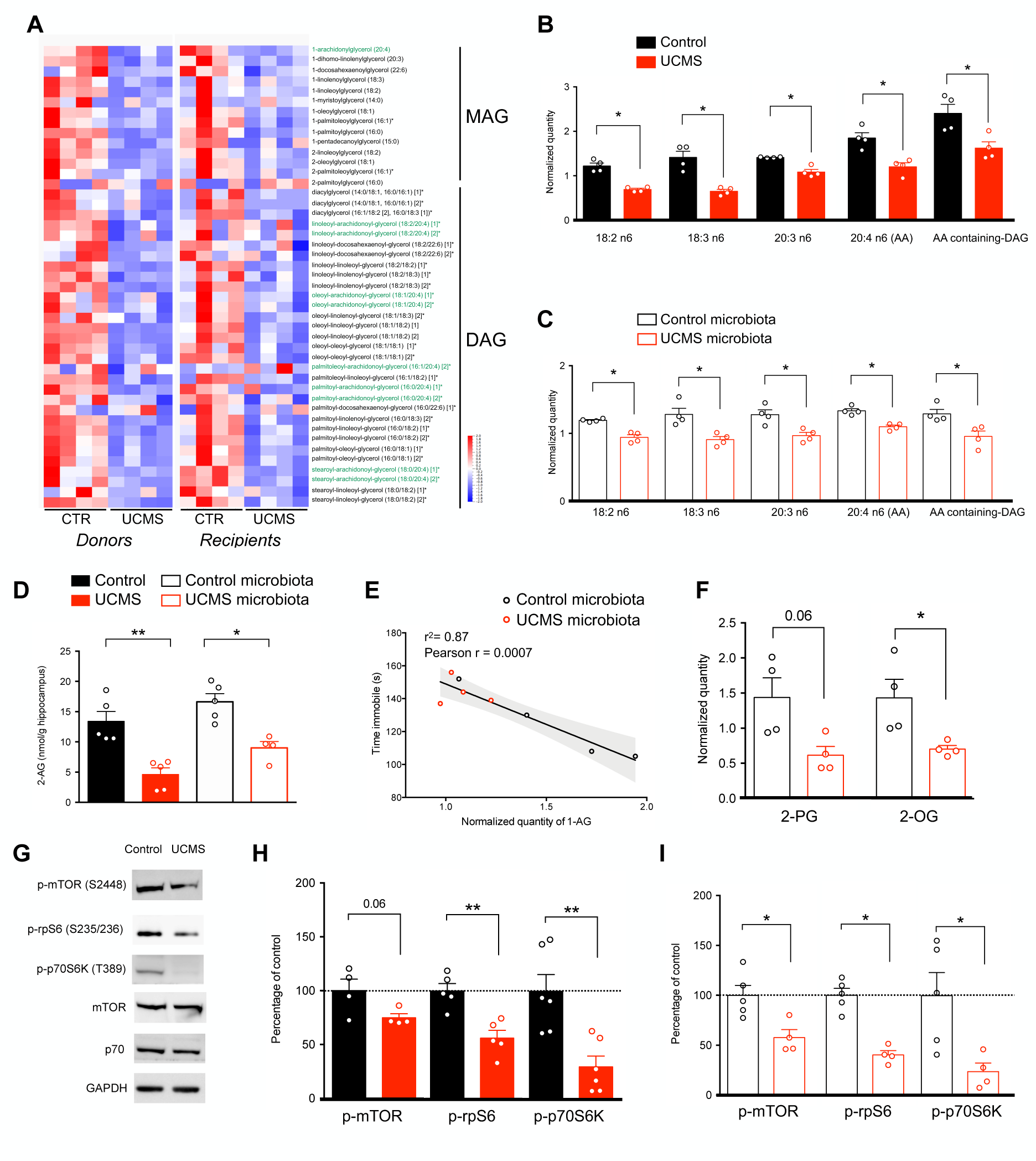
Microbiota from UCMS mice alters fatty acid metabolism and hippocampal eCB system. **A**, Heatmap of normalized serum levels of Monoacylglycerols (MAG) and Diacylglycerols (DAG) in donor (*n* = 4/group) and recipient mice (*n* = 4/group) (z-scored). Arachidonic acid-containing MAG and DAG are highlighted in green. **B-C**, Normalized levels of fatty acid in the synthesis pathway of arachidonic acid (AA) and AA-containing DAG in donor mice (*n* = 4; **B**) and in recipient mice (*n* = 4, **C**). For **B** and **C**, data are represented as mean ± s.e.m. Statistical significance was calculated using the Mann Whitney test (*, *P* = 0.0286). **D,** Concentration of 2-AG in the hippocampus of donor (*n* = 5/group) and recipient mice (*n* = 5/group) was determined by targeted LC-MS (Control vs UCMS, *P* = 0.0079; Control microbiota- vs UCMS microbiota-recipient mice, *P* = 0.0159). **E,** Correlation between serum quantity of 1-AG and time spent immobile in the tail suspension test (TST) in recipient mice (*n* = 4/group). Correlation was calculated using Pearson correlation factor r (*r* = 0.0007). **F,** Normalized quantity of the two minor endocannabinoids 2-PG and 2-OG in the serum of recipient mice (*n* = 4/group). (2-PG, *P* = 0.0571; 2-OG, *P* = 0.0286). **G,** Representative western blots for p-mTOR (S2448), p-rpS6 (S235/236), p-p70S6K (T389), mTOR, p70 and GAPDH in hippocampal protein extracts from donor mice. **H-I,** Quantification of the phosphorylation of mTOR, rpS6 and p70S6K in protein extracts from the hippocampus of Control and UCMS donor mice (p-mTOR, *n* = 4, *P* = 0.0571; p-rpS6, *n* = 5, *P* = 0.0079; p-p70S6K, *n* = 6, *P* = 0.0087; **E**) and Control microbiota-(*n* = 5) and UCMS microbiota-recipient mice (*n* = 4) (p-mTOR, *P* = 0.0317; p-rpS6, *P* = 0.0159; p-p70S6K, *P* = 0.0317; **F**). For **B**, **C, D**, **F**, **H** and **I**, data are represented as mean ± s.e.m. Statistical significance was calculated using the Mann Whitney test (\**P* < 0.05, ** *P* < 0.01).

Such changes in the levels of fatty acids could originate from altered gut permeability and/or dysbiosis-induced lipid metabolism changes. To test the first hypothesis, we quantified fluorescence level in the serum following gavage with FITC-dextran and found no change in gut permeability (**Supplementary Fig. 3E**). To address the second hypothesis, we scrutinize several fatty acid metabolites and found that two precursors for the production of eCB, AA-containing DAG and n-6 PUFA, were dramatically reduced in recipient mice transplanted with UCMS microbiota but not with control microbiota. Interestingly, dysregulation of the eCB system and its main central receptor CB1 has been associated with the pathophysiology of depression both in humans and in UCMS model of depression^44, 45^.

Since previous studies have shown that activation of the CB1 receptors produces anxiolytic and antidepressant-like effects, notably via the modulation of hippocampal neurogenesis^46–48^, we investigated into more details the brain eCB system. We examined both the hippocampal production of eCB ligands and the activation level of the CB1 receptor pathway. As the AA-containing DAG and n-6 PUFA are precursors of the endocannabinoid 2-arachidonoylglycerol (2-AG), we first compared the levels of 2-AG and its more stable metabolite 1-AG in the hippocampus and serum of donor and recipient mice^49^. Levels of hippocampal 2-AG, determined by mass spectrometry, revealed a significant decrease in both UCMS donors and recipients (Fig. 2D), with a strong inverse correlation found between the serum levels of 1-AG and the depressive state (Fig. 2E). Similar results were reported for other eCBs (Fig. 2F).

In the hippocampus, activation of CB1 receptors triggers mTOR signaling. To evaluate whether deficiency in 2-AG leads to altered activity of the mTOR pathway, we quantified phosphorylated (active) mTOR and its downstream effectors in both UCMS donor and recipient mice. mTOR phosphorylates the 70-kDa ribosomal protein S6 kinase (p70S6K) at T389^50^ and the activated p70S6K in turn phosphorylates the ribosomal protein S6 (rpS6) at S235/236, which initiates mRNA translation^51^. Donors and recipients of UCMS microbiota showed significantly decreased phosphorylation of mTOR (p-mTOR), p70S6K (p-p70S6K), and rpS6 (p-rpS6) (Fig. 2G to I). Collectively, these results demonstrate that the signature in lipid metabolism of UCMS microbiota comprises a deficiency in serum 2-AG precursors, lower content in hippocampal 2-AG and breakdown of the mTOR signaling. Remarkably, these features were found to be transmittable to naïve recipient mice following FMT.

### Restoration of eCB signaling normalizes behavior and adult neurogenesis

To further demonstrate the role of defective eCB signaling in the depressive-like behaviors of mice transferred with UCMS microbiota, we next assessed whether enhancing eCB signaling, using pharmacological blockade of the 2-AG-degrading enzyme monoacylglycerol lipase (MAGL), could alleviate these phenotypes. Recipients of UCMS microbiota were treated with the MAGL inhibitor JZL184, or JZL184 together with rimonabant, a selective antagonist of CB1, every 2 days for 4 weeks starting 4 weeks after FMT (Fig. 3A). First, we confirmed that recipients of UCMS microbiota treated with JZL184 showed a significant increase in hippocampal levels of p-mTOR, p-p70S6K, and p-rpS6 as compared to vehicle-treated recipient mice of UCMS microbiota (Fig. 3B and C). Furthermore, consistent with enhanced eCB signaling, we confirmed that JZL184 enhanced the levels of 2-AG in the hippocampus (Fig. 3D)^52, 53^. The effect of JZL184 was strictly CB1-dependent as it was reversed by the selective CB1 receptor antagonist rimonabant. As a consequence, JZL184 reduced depressive symptoms in recipients of UCMS microbiota, an effect that was blocked by rimonabant (Fig. 3E to H). To assess the relative contribution of central *versus* peripheral CB1 receptors in these depressive-like behaviors, we compare the effects of rimonabant to the effects of AM6545, a CB1 antagonist with limited brain penetrance^54^. In contrast to rimonabant, AM6545 did not reverse the antidepressant effect of JZL184 (Fig. 3G and H), indicating that central CB1 signaling is necessary to alleviate depressive-like behaviors, at least in our model.

**Figure 3.**
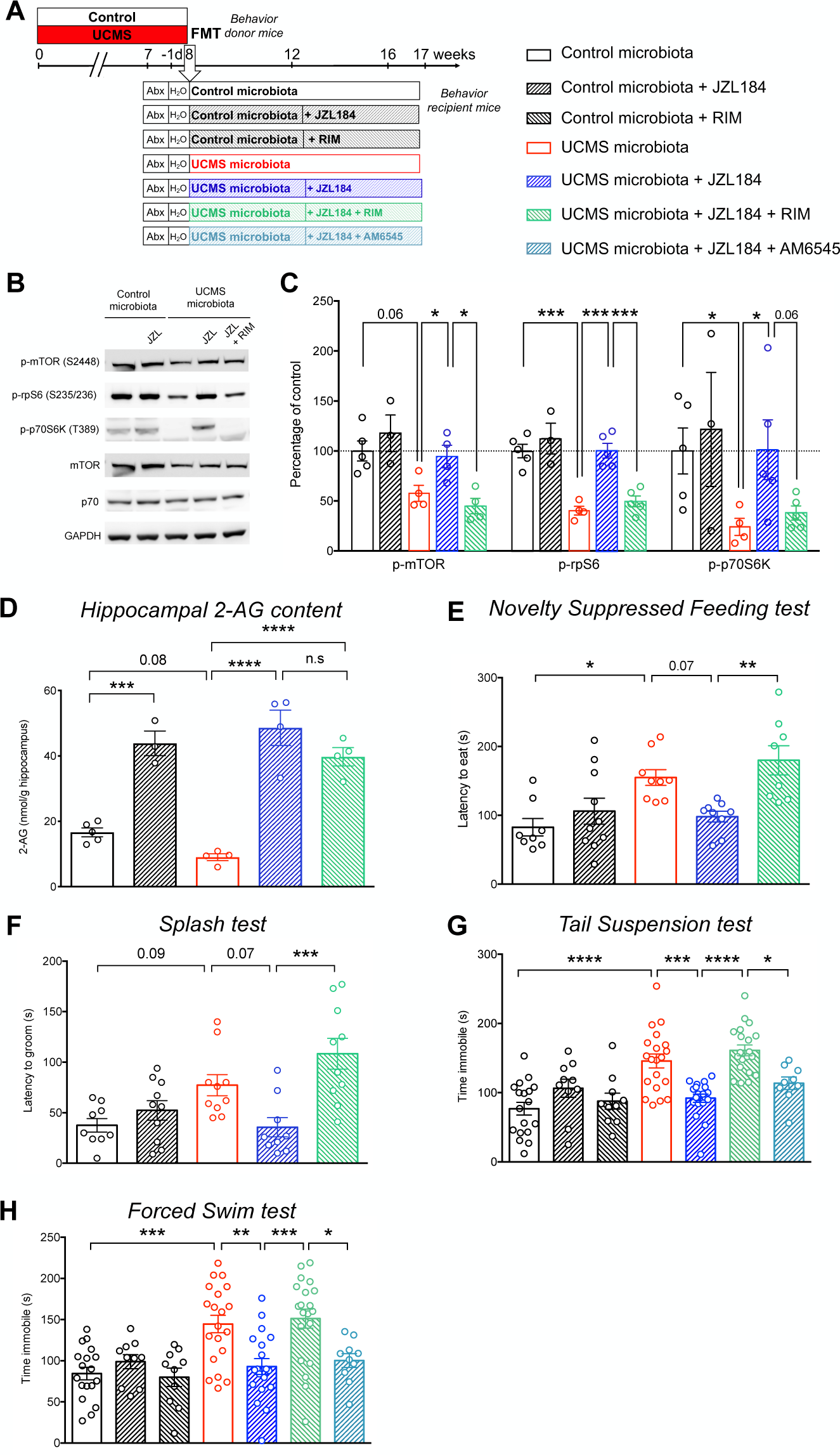
Restoration of the eCB pathway normalizes behavior in recipient mice. **A**, Experimental timeline of JZL184, rimonabant (RIM) and AM6545 treatment in recipient mice. Mice were injected intra-peritoneally every 2 days, with either vehicle alone, JZL184 (8mg/kg), rimonabant (2mg/kg), AM6545 (2mg/kg), JZL184 + rimonabant or JZL184 + AM6545. The treatment started 4 weeks after FMT and lasted for 5 weeks, until sacrifice. **B**, Representative western blots for p-mTOR (S2448), p-rpS6 (S235/236), p-p70S6K (T389), mTOR, p70 and GAPDH in hippocampal protein extracts from recipient mice upon treatment with JZL184 or rimonabant. **C,** Quantification of the phosphorylation of mTOR, rpS6 and p70S6K in hippocampal protein extracts from Control microbiota-recipient mice (*n* = 5), Control microbiota-recipient mice treated with JZL184 (*n* = 3), UCMS microbiota-recipient mice (*n* = 4), UCMS microbiota-recipient mice treated with JZL184 (*n* = 5, except for p-mTOR with *n* = 4), UCMS microbiota-recipient mice treated with JZL184 and rimonabant (*n* = 5, except for p-mTOR with *n* = 4). Control microbiota and UCMS microbiota groups are the same as in figure 2I. (p-mTOR: Control vs UCMS-recipient mice, *P* = 0.0317; UCMS-recipient mice vs UCMS-recipient mice + JZL184, *P* = 0.0571; UCMS-recipient mice + JZL184 vs UCMS-recipient mice + JZL184 + RIM, *P* = 0.0286; p-rpS6: Control vs UCMS-recipient mice, *P* = 0.0159; UCMS-recipient mice vs UCMS-recipient mice + JZL184, *P* = 0.0159; UCMS-recipient mice + JZL184 vs UCMS-recipient mice + JZL184 + RIM, *P* = 0.0079; p-p70S6K: Control vs UCMS-recipient mice, *P* = 0.0317; UCMS-recipient mice vs UCMS-recipient mice + JZL184, *P* = 0.0317; UCMS-recipient mice + JZL184 vs UCMS-recipient mice + JZL184 + RIM, *P* = 0.0556). **D**, Concentration of 2-AG in the hippocampus of Control microbiota-recipient mice (*n* = 5), Control microbiota-recipient mice treated with JZL184 (*n* = 3), UCMS microbiota-recipient mice (*n* = 4), UCMS microbiota-recipient mice treated with JZL184 (*n* = 4) and UCMS microbiota-recipient mice treated with JZL184 and rimonabant (*n* = 4), as determined by targeted LC-MS. Control microbiota and UCMS microbiota groups are the same as in figure 2D. (Control microbiota-recipient vs Control microbiota-recipient mice + JZL184, *P* = 0.0002; Control microbiota-recipient vs UCMS microbiota-recipient mice, P = 0.0872; UCMS microbiota-recipient vs UCMS microbiota-recipient mice + JZL184, *P* < 0.0001; UCMS microbiota-recipient vs UCMS microbiota-recipient mice + JZL184 + RIM, *P* < 0.0001; UCMS microbiota-recipient + JZL184 vs UCMS microbiota-recipient mice + JZL184 + RIM, *P* = 0.3037). **E**, Latency to eat in a novel environment in the Novelty Suppressed Feeding test for Control microbiota-recipient mice (*n* = 8), Control microbiota-recipient mice treated with JZL184 (*n* = 10), UCMS microbiota-recipient mice (*n* = 9), UCMS microbiota-recipient mice treated with JZL184 (*n* = 9), UCMS microbiota-recipient mice treated with JZL184 and rimonabant (*n* = 8). (Control microbiota- vs UCMS microbiota-recipient mice, *P* = 0.0179; UCMS microbiota-recipient mice vs UCMS microbiota-recipient mice + JZL184, *P* = 0.0796; UCMS microbiota-recipient mice + JZL184 vs UCMS microbiota-recipient mice + JZL184 + RIM, *P* = 0.0054). **F**, Latency to groom in the splash test for Control microbiota-recipient mice (*n* = 9), Control microbiota-recipient mice treated with JZL184 (*n* = 10), UCMS microbiota-recipient mice (*n* = 10), UCMS microbiota-recipient mice treated with JZL184 (*n* = 9), UCMS microbiota-recipient mice treated with JZL184 and rimonabant (*n* = 10). (Control microbiota- vs UCMS microbiota-recipient mice, *P* = 0.0946; UCMS microbiota-recipient mice vs UCMS microbiota-recipient mice + JZL184, *P* = 0.0721; UCMS microbiota-recipient mice + JZL184 vs UCMS microbiota-recipient mice + JZL184 + RIM, *P* = 0.0003). **G,** Time spent immobile in the Tail Suspension test for Control microbiota-recipient mice (*n* = 18), Control microbiota-recipient mice treated with JZL184 (*n* = 10), Control microbiota-recipient mice treated with rimonabant (*n* = 10), UCMS microbiota-recipient mice (*n* = 20), UCMS microbiota-recipient mice treated with JZL184 (*n* = 19), UCMS microbiota-recipient mice treated with JZL184 and rimonabant (*n* = 20) and UCMS microbiota-recipient mice treated with JZL184 and AM6545 (*n* = 9). (Control microbiota- vs UCMS microbiota-recipient mice, *P* < 0.0001; UCMS microbiota-recipient mice vs UCMS microbiota-recipient mice + JZL184, *P* = 0.0002; UCMS microbiota-recipient mice + JZL184 vs UCMS microbiota-recipient mice + JZL184 + RIM, *P* < 0.0001; UCMS microbiota-recipient mice + JZL184 + RIM vs UCMS microbiota-recipient mice + JZL184 + AM6545, *P* = 0.0266). **H,** Time spent immobile in the Forced Swim test for Control microbiota-recipient mice (*n* = 18), Control microbiota-recipient mice treated with JZL184 (*n* = 10), Control microbiota-recipient mice treated with rimonabant (*n* = 10), UCMS microbiota-recipient mice (*n* = 20), UCMS microbiota-recipient mice treated with JZL184 (*n* = 19), UCMS microbiota-recipient mice treated with JZL184 and rimonabant (*n* = 20) and UCMS microbiota-recipient mice treated with JZL184 and AM6545 (*n* = 10). (Control microbiota- vs UCMS microbiota-recipient mice, *P* = 0.0003; UCMS microbiota-recipient mice vs UCMS microbiota-recipient mice + JZL184, *P* = 0.0028; UCMS microbiota-recipient mice + JZL184 vs UCMS microbiota-recipient mice + JZL184 + RIM, *P* = 0.0004; UCMS microbiota-recipient mice + JZL184 + RIM vs UCMS microbiota-recipient mice + JZL184 + AM6545, *P* = 0.0276). Data are represented as mean ± s.e.m. For **C** to **H**, statistical significance was calculated using One-way ANOVA with Tukey’s multiple comparisons test (* *P* < 0.05, ** *P* < 0.01, *** *P* < 0.001, **** *P* < 0.0001).

JZL184 also alleviates the detrimental effects of UCMS microbiota on adult hippocampal neurogenesis. JZL184 treatment rescued the proliferation and differentiation of neural stem cells in the hippocampus of UCMS microbiota recipients, an effect that was blocked by rimonabant (Fig. 4A, B). The survival of newly-generated neurons was also increased in the hippocampus of mice treated with JZL184, and blocked by rimonabant, as shown by the quantification of newborn neurons labeled with the DNA synthesis marker EdU administered 4 weeks before analysis (Fig. 4C). According to the regions of the hippocampus, adult neurogenesis may subserve different functions: new neurons born in the dorsal hippocampus influences cognitive information processing whereas adult-born neurons of the ventral hippocampus regulate mood and stress response^55^. In the present study, the effects on UCMS microbiota on adult hippocampal neurogenesis were observed both in the dorsal and ventral regions of the hippocampus (Suppl. Fig 4). Together, these data demonstrate that the decrease in hippocampal neurogenesis and depressive-like behaviors observed in recipients of UCMS microbiota can be rescued by selectively increasing the activity of the brain eCB system.

**Figure 4.**
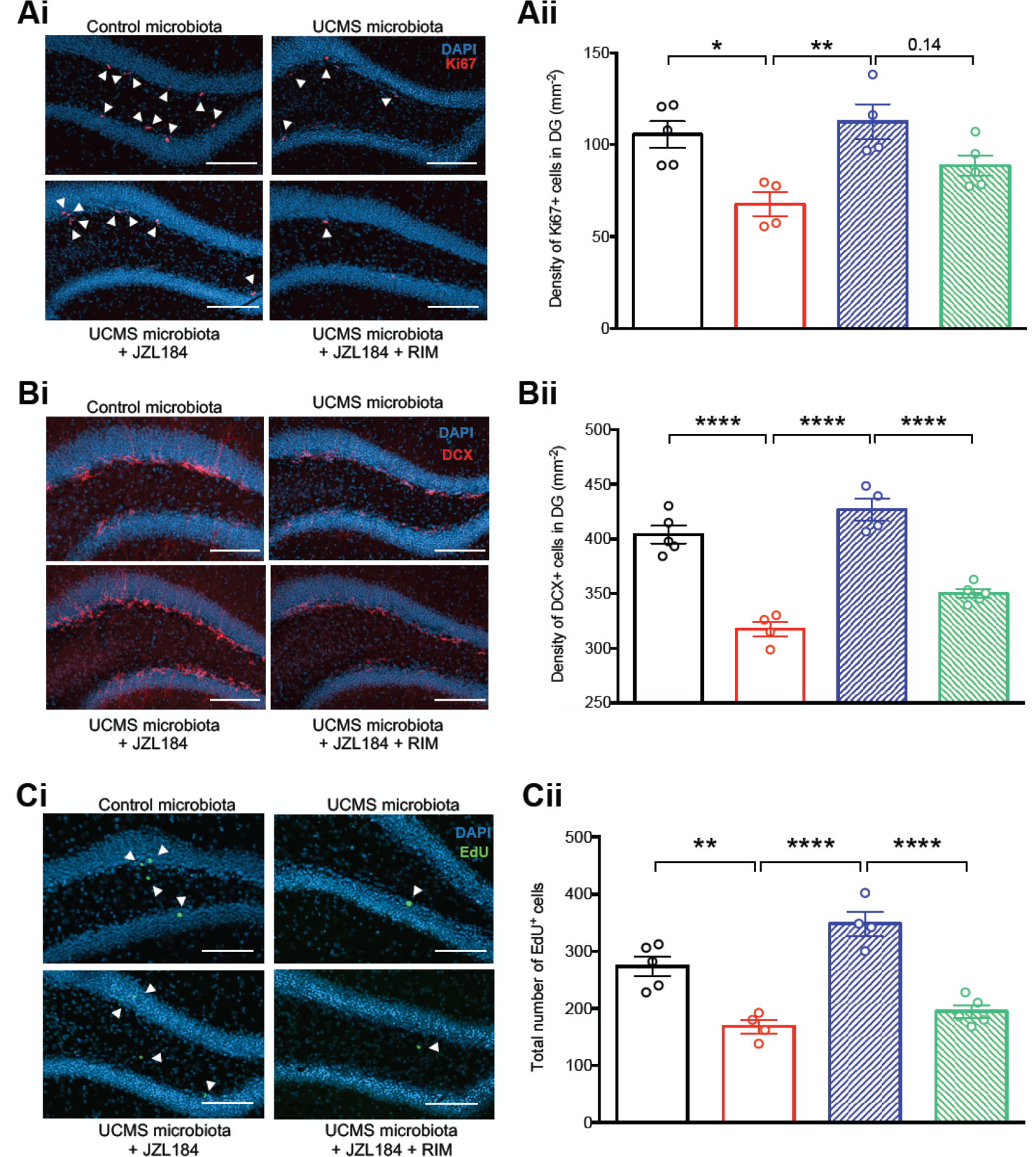
Restoration of the eCB pathway normalizes hippocampal neurogenesis. **Ai** Representative images of Ki67 staining (red) in the DG of the hippocampus, counterstained with DAPI (blue). **Aii,** Quantitative evaluation of the density of Ki6rcells for Control microbiota-recipient mice *(n* = 5), UCMS microbiota-recipient mice *(n* = 4), UCMS microbiota-recipient mice treated with JZL184 *(n* = 4), UCMS microbiota-recipient mice treated with JZL184 and rimonabant (*n* = 5). (Control microbiota- vs UCMS microbiota-recipient mice, *P* = 0.0111; UCMS microbiota-recipient mice vs UCMS microbiota-recipient mice + JZL184, *P* = 0.0048; UCMS microbiota-recipient mice + JZL184 vs UCMS microbiota-recipient mice + JZL184 + RIM, *P* = 0.1402). **Bi**, Representative images of DCX staining (red) in the DG of the hippocampus, counterstained with DAPI (blue). **Bii**, Quantitative evaluation of the density of DCX^+^ cells for Control microbiota-recipient mice (*n* = 5), UCMS microbiota-recipient mice (*n* = 4), UCMS microbiota-recipient mice treated with JZL184 (*n* = 4), UCMS microbiota-recipient mice treated with JZL184 and rimonabant (*n* = 5). (Control microbiota- vs UCMS microbiota-recipient mice, *P* < 0.0001; UCMS microbiota-recipient mice vs UCMS microbiota-recipient mice + JZL184, *P* < 0.0001; UCMS microbiota-recipient mice + JZL184 vs UCMS microbiota-recipient mice + JZL184 + RIM, *P* < 0.0001). **Ci**, Representative images of EdU staining (green) in the DG of the hippocampus, counterstained with DAPI (blue). **Cii**, Quantitative evaluation of total number of EdU^+^ cells for Control microbiota-recipient mice (*n* = 5), UCMS microbiota-recipient mice (*n* = 4), UCMS microbiota-recipient mice treated with JZL184 (*n* = 4), UCMS microbiota-recipient mice treated with JZL184 and rimonabant (*n* = 5). (Control microbiota- vs UCMS microbiota-recipient mice, *P* = 0.0014; UCMS microbiota-recipient mice vs UCMS microbiota-recipient mice + JZL184, *P* < 0.0001; UCMS microbiota-recipient mice + JZL184 vs UCMS microbiota-recipient mice + JZL184 + RIM, *P* < 0.0001). Scale bars: 100µm. Data are represented as mean ± s.e.m. Statistical significance was calculated using One-way ANOVA with Tukey’s multiple comparisons test (\**P* < 0.05, ** *P* < 0.01, **** *P* < 0.0001).

We next reasoned that if UCMS microbiota induces paucity in serum levels of eCB precursors, the complementation of diet with eCB precursors, such as arachidonic acid (AA) might normalize the levels of 2-AG and restore normal behavior. Recipient mice of UCMS microbiota were given orally AA during 5 weeks starting 3 weeks after microbiota transfer (Fig. 5A). Remarkably, we observed that AA treatment restored normal levels of hippocampal 2-AG (Fig. 5B) and reversed the depressive-like behaviors induced by UCMS microbiota (Fig. 5C and D). Furthermore, AA complementation also partially restored the production and the survival of hippocampal newborn neurons (Fig. 5E and F).

**Figure 5.**
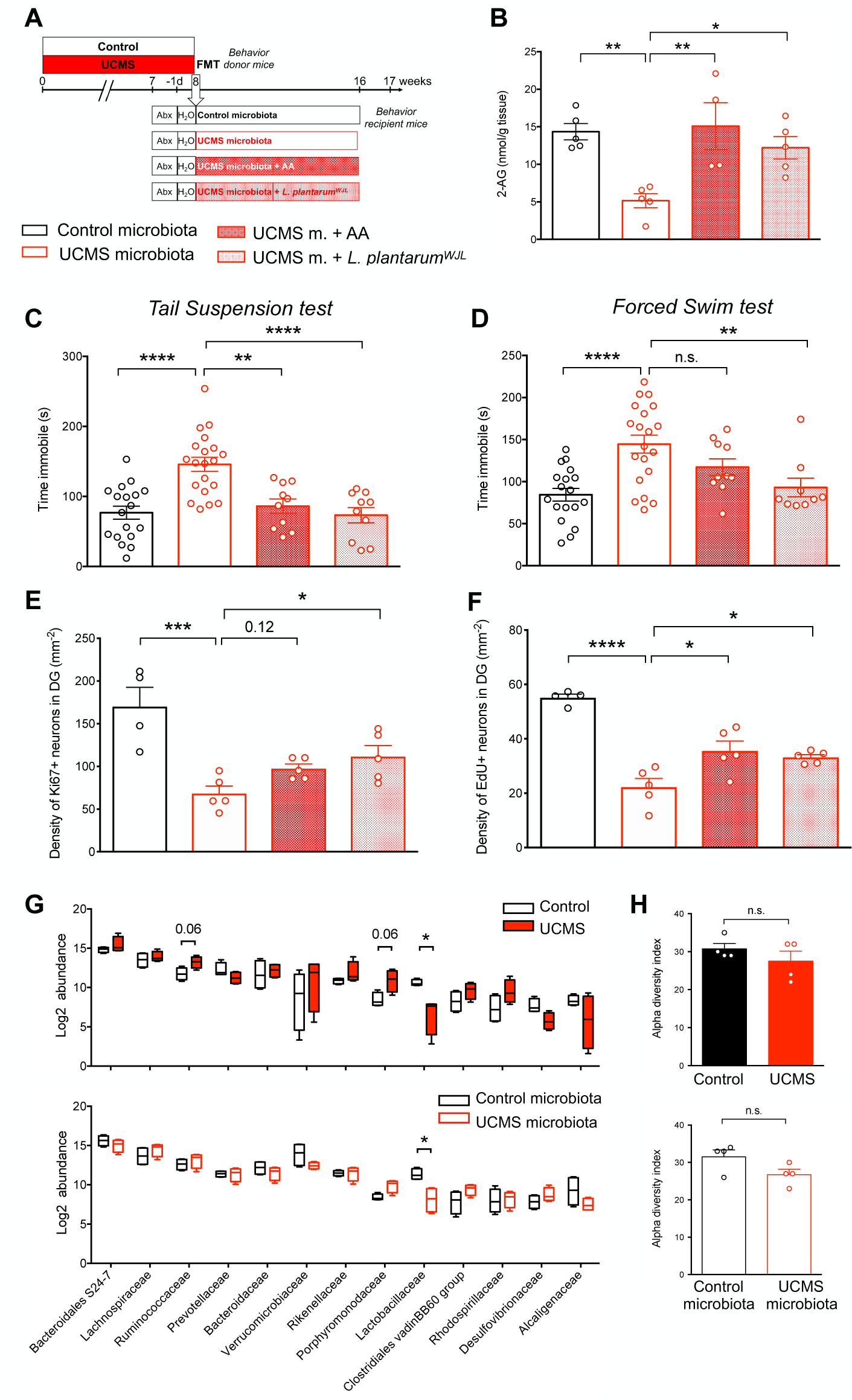
Arachidonic acid or *Lactobacillus plantarum^WJL^* complementations are sufficient to normalize hippocampal 2-AG levels, adult neurogenesis and behavior. **A**, Experimental timeline of arachidonic acid (AA) and *L. plantarum* treatment in recipient mice. Mice were fed every two days through oral gavage with 8 mg of AA/mouse/day. Mice were supplemented by oral feeding five days a week with 2×10^8^ CFU diluted in 200 μl of PBS. UCMS microbiota recipient mice were oral-fed with PBS as control. **B**, Concentration of 2-AG in the hippocampus for Control microbiota (*n* = 5), UCMS microbiota (*n* = 5), UCMS microbiota complemented with AA (*n* = 4) and UCMS microbiota complemented with *Lp^WJL^* (*n* = 5), as determined by targeted LC-MS (Control microbiota- vs UCMS microbiota-recipient mice, *P* = 0.0062; UCMS microbiota-recipient mice vs UCMS microbiota-recipient mice + AA, *P* = 0.0053; UCMS microbiota-recipient mice vs UCMS microbiota-recipient mice + *Lp^WJL^*, *P* = 0.0371). **C,** Time spent immobile in the Tail Suspension Test for Control microbiota-recipient mice (*n* = 18), UCMS microbiota-recipient mice (*n* = 20), UCMS microbiota-recipient mice complemented with AA (*n* = 10) and UCMS microbiota-recipient mice complemented with *Lp^WJL^* (*n* = 10) (Control microbiota- vs UCMS microbiota-recipient mice, *P* < 0.0001; UCMS microbiota-recipient mice vs UCMS microbiota-recipient mice + AA, *P* = 0.0015; UCMS microbiota-recipient mice vs UCMS microbiota-recipient mice + *Lp^WJL^*, *P* < 0.0001). **D,** Time spent immobile in the Forced Swim test for Control microbiota-recipient mice (*n* = 18), UCMS microbiota-recipient mice (*n* = 20), UCMS microbiota-recipient mice complemented with AA (*n* = 10) and UCMS microbiota-recipient mice complemented with *Lp^WJL^* (*n* = 9) (Control microbiota- vs UCMS microbiota-recipient mice, *P* < 0.0001; UCMS microbiota-recipient mice vs UCMS microbiota-recipient mice + AA, *P* = 0.2690; UCMS microbiota-recipient mice vs UCMS microbiota-recipient mice + *Lp^WJL^*, *P* = 0.0083). Data are represented as mean ± s.e.m. Statistical significance was calculated using one-way ANOVA with Tukey’s multiple comparisons test (** *P* < 0.01, **** *P* < 0.0001). **E**, quantitative evaluation of the density of Ki67^+^ cells for Control microbiota-recipient mice (*n* = 4), UCMS microbiota-recipient mice (*n* = 5), UCMS microbiota-recipient mice complemented with AA (*n* = 5), UCMS microbiota-recipient mice complemented with *Lp^WJL^* (*n* = 5) (Control microbiota- vs UCMS microbiota-recipient mice, *P=* 0.0003; UCMS microbiota-recipient mice vs UCMS microbiota-recipient mice + AA, *P* = 0.1175; UCMS microbiota-recipient mice vs UCMS microbiota-recipient mice + *Lp^WJL^*, *P* = 0.0258). **F**, Quantitative evaluation of the density of EdU^+^ cells for Control microbiota-recipient mice (*n* = 4), UCMS microbiota-recipient mice (*n* = 5), UCMS microbiota-recipient mice complemented with AA (*n* = 5), UCMS microbiota-recipient mice complemented with *Lp^WJL^* (*n* = 5) (Control microbiota- vs UCMS microbiota-recipient mice, *P* < 0.0001; UCMS microbiota-recipient mice vs UCMS microbiota-recipient mice + AA, *P* = 0.0113; UCMS microbiota-recipient mice vs UCMS microbiota-recipient mice + *Lp^WJL^*, *P* = 0.0394). **G**, 16S rDNA of the fecal microbiota of donor mice at the end of the 8 weeks UCMS protocol (*n* = 4/group, top) or recipients mice after 8 weeks in isolators (*n* = 4/group, bottom), was sequenced and analyzed by principal Component Analysis (PCA) at the level of bacterial families for the relative abundance of bacterial families. Data are represented as boxplots. Statistical significance was calculated using Mann-Whitney test (top, *Ruminococcaceae*, *P* = 0.0571; *Porphyromonadaceae*, *P* = 0.0571; *Lactobacillaceae*, *P* = 0.0286; bottom, *Lactobacillaceae*, *P* = 0.0286). **H**, Alpha diversity for donors (*P* = 0.6857, top) and recipients (*P* = 0.2286). Data are represented as mean ± s.e.m. Statistical significance was calculated using Mann-Whitney test. For **B** to **F**, data are represented as mean ± s.e.m. Statistical significance was calculated using one-way ANOVA with Tukey’s multiple comparisons test (* *P* < 0.05, *** *P* < 0.0005, **** *P* < 0.0001).

### UCMS-induced dysbiosis and complementation with *Lactobacillus plantarum^WJL^*

We next investigated how UCMS affected the composition of the microbiota that was responsible for the observed cellular and behavioral impairments in recipient mice. The composition of the fecal microbiota was determined by sequencing of 16S rDNA. Analysis of bacterial families revealed significant modifications in the microbiota of UCMS mice, as compared to the microbiota of control mice raised in separate cages (Fig. 5G), while the total number of species (alpha diversity) did not vary significantly (Fig. 5H). In-depth analysis of bacterial families showed an increase in *Ruminococcaceae*, and *Porphyromonodaceae*, as well as a decrease in *Lactobacillaceae* in UCMS mice (Fig. 5G and **Supplementary Fig. 5**). These results are in agreement with recent studies reporting an association between low frequencies of *Lactobacilli* and stress in mice^56–58^ or depression in patients^59^. Importantly, the differences in microbiota composition between recipient mice of UCMS and control microbiota were maintained 8 weeks after transfer (Fig. 5G), in particular the decrease in *Lactobacillaceae* (Fig. 5G and **Supplementary Fig. 5**), while the total number of species (alpha diversity) did not vary (Fig. 5H).

Since the frequencies of *Lactobacillaceae* were decreased in UCMS microbiota when compared to control microbiota (Fig. 5G), we tested whether complementation of UCMS microbiota with *Lactobacillaceae* normalized behaviors and neurogenesis levels in recipients of UCMS microbiota. To this end, the microbiota of recipients was complemented with a strain of *Lactobacillus plantarum* (*Lp^WJL^*) shown to modulate the host’s lipid composition^60, 61^, to stimulate juvenile growth^62^ and to influence affective behavior in mice^63^. Recipient mice of UCMS microbiota were given orally *Lp^WJL^* for 5 weeks starting 3 weeks after microbiota transfer (Fig. 5A). We observed that *Lp^WJL^* restored normal levels of hippocampal 2-AG (Fig. 5B), reversed the depressive-like behaviors induced by UCMS microbiota (Fig. 5C and D) and partially restored the production and the survival of hippocampal newborn neurons (Fig. 5E and F). These data indicate that *Lactobacillaceae* play an important role in the host metabolism with significant effect on mood control.

## DISCUSSION

In the present study, we have explored the mechanisms by which gut microbiota dysbiosis contributes to brain dysfunctions and behavioral abnormalities associated with depressive-like states. Chronic stress is recognized as a major risk factor for depression^64^ and most animal models of depressive-like behaviors rely on chronic stress or manipulation of the stress-sensitive brain circuits^65^. Using UCMS as a mouse model of depression, we showed that, upon transplantation to naïve hosts, the microbiota from UCMS mice reduced adult hippocampal neurogenesis and induced depressive-like behaviors.

Searching for mechanistic explanations of these dysfunctions, we found that UCMS microbiota alters the fatty acid metabolism of the host, leading to paucity in precursors of the eCB system, such as AA, reduced production of the eCB 2-AG in the hippocampus, and diminished signaling in the hippocampal eCB system. Restoration of normal eCB signaling levels in mice recipient of UCMS microbiota after blocking the 2-AG-degrading enzyme, or after complementation of the diet with the 2-AG precursor AA, both restored adult neurogenesis and behaviors. Finally, UCMS-induced perturbations of the gut bacterial composition were characterized by loss of *Lactobacillaceae,* an alteration that was maintained after microbiota transplantation to naïve hosts. The mere complementation of the UCMS recipients’ microbiota with *Lactobacillus plantarum Lp^WJL^* was sufficient to normalize the levels of 2-AG in the hippocampus and restore affective behaviors and adult hippocampal neurogenesis.

The eCB system has been reported to regulate mood, emotions and responses to stress through activation of the cannabinoid receptor CB1. For instance, the CB1 receptor antagonist rimonabant, initially prescribed for the treatment of obesity and associated metabolic disorders, increases the incidence of depressive symptoms^66^. Furthermore, a higher frequency in a mutant allele for the CB1 receptor gene *CNR1* is observed in depressed patients^67^. In contrast, cannabis (that includes the eCB ligand delta-9 tetrahydrocannabinol or THC) improves mood in humans^68^ and synthetic CB1 agonists produce anxiolytic- and antidepressant-like effects in animal models^69^. In particular, chronic stress has been showed to decrease eCB signaling in the brain^70–74^. Here, we show that the intestinal microbiota is sufficient to initiate a pathological feed-forward loop for depressive disorders by impairing the eCB system in the hippocampus, a brain region strongly involved in the development of depressive symptoms. Previous studies have shown that the reduction in hippocampal CB1 signaling, involving mTOR, induces depressive-like behaviors^75^, and different studies on postmortem brains of depressed patients have shown deficits in mTOR signaling^76, 77^. In line with this, we observed a specific decrease in 2-AG, one of the two major eCB ligands, in mice receiving UCMS-derived microbiota, but not medium- or long-chain fatty acids. This result is reminiscent to clinical observations reporting low serum levels of 2-AG in patients suffering from depression, post-traumatic stress disorder or chronic stress, but not of the other main eCB ligand anandamide^78–80^.

The eCB system exerts its pleiotropic effects through multiples neuronal processes including, but not limited to, adult hippocampal neurogenesis. The eCB system is known to regulates adult neurogenesis via the CB1 receptor^81^ expressed by neural progenitor cells^46, 48^. CB1-deficient mice show impaired neural progenitor proliferation, self-renewal, and neurosphere generation^46^ whereas CB1 receptor agonists increase neurogenesis^69, 82, 83^. In addition to this neurogenic effect occurring in the hippocampus, other CB1 receptor-dependent processes might contribute to the pathophysiology of our microbiota-induced depression. Further studies should be conducted to test whether other brain targets of eCB signaling are equally affected by microbiota dysbiosis.

It has been reported previously that the microbiota modulates the activity of the eCB system in the gut^84–87^. In the present study, we further demonstrate that the dysbiotic gut microbiota from UCMS mice is sufficient to induce dysregulation of the eCB system in the brain. We report that this dysregulation originates from a systemic decrease in eCB precursors. Modifications in gut microbiota composition following chronic stress has been extensively reported^56, 58, 88–90^. In particular, low frequencies of *Lactobacillaceae* are correlated with stress levels in mouse models^56–58^. Dysbiosis of the gut microbiota and low *Lactobacilli* frequency have also been detected in depressed patients^20, 21, 28, 59^, and transplantation of the microbiota from these patients into germ-free mice induces depressive- or anxiety-like behaviors in the recipients^28, 91^. In line with these results, a probiotic treatment with *Lactobacilli* ameliorates depressive- and anxiety-like behaviors in mice^58, 63^. Gut microbiota also modulates adult neurogenesis^92, 93^ and a *Lactobacillus* strain has been shown to promote the survival of hippocampal neuronal progenitor^92^. Numerous studies have shown that *Lactobacilli* treatment, as well as the administration of other probiotics, are beneficial in significantly lowering depression and anxiety scores in patients^94–100^. *L. plantarum* in particular was recently shown to alleviate stress and anxiety^101^. Our study demonstrating the beneficial effects of *L. plantarum^WJL^* to complement a maladaptive microbiota adds to several emerging evidence showing an antidepressant effects of probiotics in major depression^99, 100^. We have found that one of the mechanisms by which *Lactobacilli* promotes these effects is through regulation of the bioavailability of eCB precursors.

A major finding of our study is that recipients of UCMS microbiota developed an altered fatty acid metabolism characterized by deficiency in MAG, DAG and fatty acids. Serum levels of MAG, DAG and PUFAs were inversely correlated with the severity of depressive-like behaviors. Further studies should clarify whether serum levels of fatty acid could be considered as earlier biomarker for mood disorders. It has been reported that nutritional n-3 PUFA deficiency abolishes eCB-mediated neuronal functions^102^, and conversely, that n-3 PUFA dietary supplementation reverses some aspects of UCMS-induced depressive-like behaviors in mice^102, 103^. We may speculate that UCMS microbiota promotes the degradation of PUFA or alters the absorption of these fatty acids. The mechanisms by which gut microbiota modulates the host’s fatty acid metabolism has been partially investigated in several animal models. Microbiota regulates intestinal absorption and metabolism of fatty acids in the zebrafish^104, 105^, and in rodents, *Lactobacilli* species modulate lipid metabolism^106, 107^. Specifically, *Lactobacillus plantarum* modulates the host’s lipid composition by reducing the level of serum triglycerides in the context of high fat diet^60, 108, 109^. Furthermore, in humans, *Lactobacillus plantarum* is associated with lower levels of cholesterol^61^. It is proposed that *Lactobacillus plantarum* regulates fatty acid metabolism and modifies fatty acid composition of the host^107^.

In sum, our data show that microbiota dysbiosis induced by chronic stress affects lipid metabolism and the generation of eCBs, leading to decreased signaling in the eCB system and reduced adult neurogenesis in the hippocampus. This might be the pathway, at least in part, that links microbiota dysbiosis to mood disorders, which in turn, may affect the composition of the gut microbiota through physiological adjustments and modulation of the immune system. Because we were able to interrupt this pathological feed-forward loop by administrating arachidonic acid or a *Lactobacillus* probiotic strain, our study supports the concept that dietary or probiotic interventions might be efficient weapons in the therapeutic arsenal to fight stress-associated depressive syndromes.

## ACKOWLEDGEMENTS

We would like to thank all the members of the Eberl and Lledo lab, as well as members of the Peduto lab for insightful discussions, and Pr. Peduto for access to the Apotome microscope. A special thanks also to the members of the Pasteur Animal Facility who were essential for this project, and in particular Marion Bérard, Martine Jacob, Thierry Angelique and Eddie Maranghi. This work was supported by the Major Federating Program “Microbes & Brain” of the Institut Pasteur, Agence Nationale de la Recherche Grant ANR-16-CE15-0021-02-PG-Brain, and a grant from the Fédération pour la Recherche sur le Cerveau (FRC). The Lledo’s lab is supported by Agence Nationale de la Recherche Grants ANR-16-CE37-0010-ORUPS and ANR-15-NEUC-0004-02 “Circuit-OPL,” Laboratory for Excellence Programme “Revive” Grant ANR-10-LABX-73, and the Life Insurance Company AG2R-La Mondiale.

## AUTHOR CONTRIBUTIONS

G.C., G.E. and P.M.L. conceived the study; G.C. established the methodology; G.C., E.S., M.P., L.G.M., T.L. and A.R. performed the experiments; G.C. wrote the original manuscript, which was edited by all authors; G.C., G.E., P.M.L., I.G.B. and G.L. secured funds; G.E., P.M.L., I.G.B. and C.D. provided resources, and G.E., P.M.L, I.G.B. and C.D. supervised the project.

## MATERIALS AND METHODS

### Mice

Adult male C57BL/6J mice (8-10 weeks old) were purchased from Janvier laboratories (St Berthevin, France) and maintained under specific-pathogen free (SPF) conditions at the Institut Pasteur animal care facility. Germ-free C57BL/6J mice were generated at the Gnotobiology Platform of the Institut Pasteur and routinely monitored for sterility. Mice were provided with food and water *ad libitum* and housed under a strict 12 h light-dark cycle. All animal experiments were approved by the committee on animal experimentation of the Institut Pasteur and by the French Ministry of Research.

### Fecal Microbiota Transplantation (FMT) Protocol

Recipient mice were given a combination of vancomycin (0.5 g/l), ampicillin (1 g/l), streptomycin (5 g/L), colistin (1 g/l), and metronidazole (0.5 g/l) in their drinking water for 6 consecutive days. All antibiotics were obtained from Sigma Aldrich (St Quentin Fallavier, France). Twenty-four hours later, animals were colonized via two rounds of oral gavage with microbiota, separated 3 days apart, and kept in separate sterile isolators. Donor microbiota was acquired from pooled fecal samples from 5-6 animals and resuspended in PBS.

### Unpredictable Chronic Mild Stress (UCMS) Protocol

After one week of habituation to the Institut Pasteur facility upon arrival, mice were subjected to various and repeated unpredictable stressors several times a day during 8 weeks. During exposure to stressors, mice of the UCMS group were housed in a separate room. The stressors included altered cage bedding (recurrent change of bedding, wet bedding, no bedding), cage tilting (45°), foreign odor (new cage impregnated with foreign mouse urine), restraint (1h-1h30 in a clean 50 mL conical tube with pierced holes for ventilation), altered light/dark cycle. On average, two stressors were administered per day. The timeline of the stressor exposure is described in **Supplementary Table S1**. For stressed animals, cages were changed after ‘wet bedding’ and ‘no bedding’ stressors. Unstressed controls were handled only for injections, cage changes and behavioral tests.

### CB1 Antagonists and JZL184 Treatment

JZL184, rimonabant and AM6545 were purchased from Cayman Chemicals (Bertin Technologies, Montigny-le-Bretonneux, France). The drugs were dissolved in a vehicle containing a 1:1:18 mixture of ethanol, kolliphor, and saline, and injected intra-peritoneally (i.p.) at a volume of 10 μl.g^−1^ bodyweight every 2 days. Mice were injected with either vehicle alone, JZL184 (8mg/kg), rimonabant (2mg/kg), AM6545 (2mg/kg), JZL184 + rimonabant or JZL184 + AM6545. The dose and treatment time of drug administration, alone or in combination, were chosen based on previous studies showing that JZL184 irreversibly inhibits the monoacylglycerol lipase (MAGL) and produces at least two-fold increase in 2-arachidonoylglycerol (2-AG) levels in the brain at a dose of 8 mg/kg when dissolved in the vehicle used in this study^1, 2^. Repeated administration of JZL184 at this low dose does not induce observable CB1 receptor desensitization or functional tolerance^3^.

### Arachidonic Acid and Lactobacilli Complementation

Arachidonic acid (AA) was purchased from Cayman Chemicals (Bertin Technologies). Mice were fed every two days through oral feeding gavage with 8 mg of AA/mouse/day. *Lactobacillus plantarum Lp^WJL^* was kindly provided by Pr. François Leulier (ENS, Lyon, France) and mice were supplemented by oral feeding five days a week with 2×10^8^ CFU diluted in 200 μl of PBS. UCMS microbiota recipient mice were free-fed with only PBS as control.

### Microbial DNA Extraction and 16S Sequencing

Total DNA was extracted from feces using the FastDNA Spin kit, following the instructions of the manufacturer (MP Biomedicals). DNA concentrations were determined by spectrophotometry using a Nanodrop (Thermo Scientific). Microbial composition was assessed by 16S metagenomic analysis, performed on an Illumina MiSeq instrument using a v3 reagent kit. Libraries were prepared by following the Illumina “16S Metagenomic Sequencing Library Preparation” protocol (Part # 15044223 Rev. B) with the following primers: Forward-5’-TCGTCGGCAGCGTCAGA TGTGTATAAGAGACAGCCTACGGGNGG-CWGCAG-3’; Reverse-5’-GTCTCGTGG GCTCGGAGATGTGTATAAGAGACAGGA-CTACHVGGGTATCTAATCC-3’. PCR amplification targeted the V3-V4 region of the 16s rDNA. Following purification, a second PCR amplification was performed to barcode samples with the Nextera XT Index Primers. Libraries were loaded onto a MiSeq instrument and sequencing was performed to generate 2 x 300 bp paired-ends reads. De-multiplexing of the sequencing samples was performed on the MiSeq and individual FASTQ files recovered for analysis.

### 16S Data Analysis

Sequences were clustered into OTUs (Operational Taxonomic Units) and annotated with the MASQUE pipeline (https://github.com/aghozlane/masque) as described^4^. OTU representative sequences were assigned to the different taxonomic levels using RDP Seqmatch (RDP database, release 11, update 1)^5^. Relative abundance of each OTU and other taxonomic levels was calculated for each sample in order to consider different sampling levels across multiple individuals. After trimming, numbers of sequences clustered within each OTU (or other taxonomic levels) were converted to relative abundances. Statistical analyses were performed with SHAMAN (shaman.c3bi.pasteur.fr) as described^6^. Briefly, the normalization of OTU counts was performed at the OTU level using the DESeq2 normalization method. In SHAMAN, a generalized linear model (GLM) was fitted and vectors of contrasts were defined to determine the significance in abundance variation between sample types. The resulting *P-*values were adjusted for multiple testing according to the Benjamini and Hochberg procedure^7^. Principal coordinates analysis (PCoA) was performed with the *ade4* R package (v.1.7.6) using a Bray-Curtis dissimilarity matrix. Further statistical analysis was conducted using Prism software (GraphPad, v6, San Diego, USA).

### Gut permeability test

This examination is based on the intestinal permeability to 4kD fluorescent-dextran (Sigma-Aldrich). After 4 hours of food withdrawal, mice were orally administered with FITC-dextran (0,6 g/kg body weight). After 1 hour, 200μl of blood was collected in Microvette^®^ tube (Sarstedt, Marnay, France). The tubes were then centrifuged at 10 000g for 5 minutes, at room temperature, to extract the serum. Collected sera were diluted with same volume of PBS and analyzed for FITC concentration at excitation wavelength of 485 nm and the emission wavelength of 535 nm.

### Behavioral Assays

Anxiety and depressive-like behaviors were assessed at time points of interest. Mice were tested for light/dark box, splash test, novelty suppressed feeding, tail suspension test and forced swim test, in that order. In order to limit the eventual microbiota divergence once the recipient mice were removed from the isolators, behavioral tests were performed within a week, with at least 24 hours between each behavioral test. Order of passage between groups was randomized.

Anxiety-like behaviors were evaluated in the light/dark box (LDB) tests. Depressive-like behaviors were evaluated in the splash test, the novelty suppressed feeding test, the tail suspension test and the forced swim test.

- *Light/Dark (L/D) Box*. The test was conducted in a 44×21×21 cm Plexiglas box divided into dark and light compartments separated by an open door. The light in the light compartment was set up at 300 lux. Time spent in the light compartment and transitions between compartments during 10 min were video-tracked using EthoVision XT 5.1 software (Noldus Information Technology).
- *Splash test.* The splash test consists of squirting a 10% sucrose solution on the dorsal coat of a mouse in its home cage. Because of its viscosity, the sucrose solution dirties the mouse fur and animals initiate grooming behavior. After applying sucrose solution, latency to grooming, frequency and time spent grooming was recorded for a period of 6 minutes as an index of self-care and motivational behavior. The splash test, pharmacologically validated, demonstrates that UCMS decreases grooming behavior, a form of motivational behavior considered to parallel with some symptoms of depression such as apathetic behavior^8–10^.
- *Novelty Suppressed Feeding (NSF).* The NSF was carried out similar to a published protocol^9^. Mice were deprived of food for 24h before being placed in a novel environment, a white plastic box (50×50×20cm) whose floor was covered with wooden bedding. A single food pellet (regular chow) was placed on a piece of filter paper (10cm in diameter), positioned in the center of the container that was brightly illuminated (∼500 lux). The mouse was placed in one corner of the box and the latency to feed was measured during 10 min. Feeding was defined as biting not simply sniffing or touching the food. Immediately after the test, the animals were transferred into their home cage and the amount of food consumed over the subsequent 5 min period were measured as a control of feeding drive.
- *Tail Suspension test.* Mice were suspended by the tail using adhesive tape affixed 1cm from the origin of the tail, on a metal rod under dim light conditions (∼40 lux). The behavior of the animals was recorded by a video camera during a 5 min period and total immobility time was evaluated in a blind manner.
- *Forced Swim test.* Mice were placed individually into plastic cylinders (19cm diameter, 25cm deep) filled to a depth of 18 cm with water (23-25°C) under dim light conditions (∼40 lux) for 5 min. The behavior of the animals was recorded by a video camera and immobility time was automatically evaluated using EthoVision XT 5.1 software (Noldus Information Technology). In both TST and FST, mice face an uncomfortable situation that they confront by attempting to move out of it, and eventually surrender to.

### 5-Ethynyl-2’-deoxyuridine (EdU) Labeling

The study of proliferation and differentiation of neural stem cells in the dentate gyrus was performed by incorporation of 5-ethynyl-2’-deoxyuridine (EdU, Click-iT EdU Imaging Kit; Molecular Probes) to allow the analysis of proliferation and differentiation. Mice received four intraperitoneal injections (100 mg/kg), at 2 h intervals, on a single day, 4 weeks before perfusion, for the analysis of cell survival. EdU incorporation was visualized as described in the immunohistochemistry section.

### Immunohistochemical Analysis

Mice were deeply anesthetized with sodium pentobarbital (i.p., 100 mg/ kg, Sanofi) and perfused transcardially with a solution containing 0.9% NaCl and heparin (Sanofi-Synthelabo), followed by 4% paraformaldehyde in phosphate buffer, pH 7.3. Brains were removed and postfixed by incubation in the same fixative at 4°C overnight. Tissues were cryoprotected by incubation in 30% sucrose in PBS for 24 h. Immunostaining was performed on 40-µm or 60-µm thick coronal brain sections obtained with a vibrating microtome (VT1000S, Leica). Nonspecific staining was blocked by 0.2% Triton, 4% bovine serum albumin (Sigma-Aldrich) and 2% goat serum and free-floating slices were then incubated with the following primary antibodies at 4°C overnight: rabbit anti-DCX (Abcam, ab 18723), rabbit anti-Ki67 (Abcam, ab16667), mouse anti-NeuN (Millipore, MAB377). Secondary antibodies (Alexa Fluor-conjugated secondary antibodies, Molecular Probes) were then incubated at room temperature. DAPI (1µg/mL) was used as a nuclear stain. EdU was visualized using the Click-iT reaction coupled to an Alexa Fluor® azide following the instructions of the manufacturer (Molecular Probes).

### Image Acquisition and Quantification Analysis

Immunofluorescence was analyzed using an Apotome microscope (Apotome.2; Zeiss) with Zen Imaging software (Zeiss), courtesy of Pr. Peduto. Quantification was performed using the Icy open source platform (http://www.icy.bioimageanalysis.org)^11^. The region of interest was defined as the granule cell layer (GCL) of the dentate gyrus and automatic detection of Ki67^+^ and DCX^+^ cells was performed using the spot detector tool. Values are expressed as the mean of total Ki67^+^ or DCX^+^ cell count per mm^2^ in six slices per animal. All imaging and quantification were performed blinded to experimental conditions. For EdU analysis, positive cells were manually counted in the GCL of the DG. Total number was estimated by multiplying the total number of cells every sixth section by six.

### Western Blotting

Mice were deeply anesthetized with sodium pentobarbital (i.p.100 mg/kg, Sanofi) and rapidly decapitated. The hippocampi were bilaterally dissected out and then homogenized in 0.2 ml lysis buffer (pH 7.5) containing 20 mM Tris-acetate, 150mM NaCl, 50 mM NaF, 1 mM EDTA, 1% Triton-X100, 0.1% benzonase, protease inhibitors and protein phosphatase inhibitors I and II (Sigma-Aldrich). After an incubation of 30 min on ice and centrifugation at 10 000g for 10 min, total protein concentration of the supernatant was assayed by using Bio-Rad protein assay kit (Bio-Rad, Marnes-la-Coquette, France). Equal amounts of each protein sample were separated on NuPAGE Bis-Tris or Tris-Acetate gels and transferred to nitrocellulose or PVDF membranes, respectively. Blots were blocked in blocking buffer containing 5% (w/v) milk and 0.1% (v/v) Tween-20 in Tris-buffered saline (TBS-T) for 1∼2 hours at room temperature, and incubated overnight at 4°C with antibodies against p-mTOR (S2448) (1:1000, Cell Signaling), mTOR (1:1000, Cell Signaling), p-p70S6K (T389) (1:500, R&D Systems), p-rpS6 (S235/236) (1:500, R&D Systems) or GAPDH (1:1000, Cell Signaling) antibodies. Blots were washed 3 times with TBS-T and then probed with anti-rabbit IgG, HRP-linked antibody (1:3000, Cell Signaling) for 1 hour at room temperature before being revealed using ECL Prime detection reagent (GE Healthcare) and chemiluminescence reading on a luminescent image analyzer (LAS-4000; Fujifilm). Immunoreactivity of Western blots was quantified by densitometry using the ImageJ software (NIH, Bethesda).

### Biochemical Detection of 2-AG

Mice were deeply anesthetized with sodium pentobarbital (i.p.100 mg/ kg, Sanofi) and decapitated. The brain was immediately removed, and the hippocampi were dissected out and rapidly frozen on dry ice. 2-AG was extracted from the hippocampus as previously described^12^. Samples were weighed and placed into borosilicate glass culture tubes containing 2 ml of acetonitrile with 186 pmol [^2^H_8_] 2-AG. They were homogenized using IKA homogenizer and kept overnight at -20°C to precipitate proteins and subsequently centrifuged at 1500g for 3 min. The supernatants were transferred to a new glass tube and evaporated to dryness under N2 gas. The samples were resuspended in 500µl of methanol to recapture any lipids adhering to the glass tube and dried again under N2 gas. Dried lipid extracts were suspended in 50µl of methanol and stored at -80°C until analysis. The content of 2-AG was determined using isotope-dilution liquid chromatography–electrospray ionization tandem mass spectrometry (LC-MS/MS)^13^.

### Metabolomics

Blood were collected by cardiac puncture in Microvette^®^ tubes (Sarstedt, Marnay, France), from behaviorally validated adult mice. The tubes were centrifuged at 10 000g for 5 minutes, at room temperature, to extract the serum. Serum samples were then extracted and analyzed on GC/MS, LC/MS and LC/MS/MS platforms by Metabolon, Inc (California, USA). Protein fractions were removed by serial extractions with organic aqueous solvents, concentrated using a TurboVap system (Zymark) and vacuum dried. For LC/MS and LC/MS/MS, samples were reconstituted in acidic or basic LC-compatible solvents containing > 11 injection standards and run on a Waters ACQUITY UPLC and Thermo-Finnigan LTQ mass spectrometer, with a linear ion-trap front-end and a Fourier transform ion cyclotron resonance mass spectrometer back-end. For GC/MS, samples were derivatized under dried nitrogen using bistrimethyl-silyl-trifluoroacetamide and analyzed on a Thermo-Finnigan Trace DSQ fast-scanning single-quadrupole mass spectrometer using electron impact ionization. Chemical entities were identified by comparison to metabolomic library entries of purified standards. Following log transformation and imputation with minimum observed values for each compound, data were analyzed using two-way ANOVA with contrasts.

### Statistical analysis

Statistical analysis was performed using Prism software (GraphPad, v6, San Diego, USA). Principal component analyses (PCA) and heatmaps were performed using Qlucore Omics Explorer (Qlucore). Data are plotted in the figures as mean ± SEM. Differences between two groups were assessed using Mann-Whitney test. Differences among three or more groups were assessed using one-way ANOVA with Tukey’s. Significant differences are indicated in the figures by *p < 0.05, **p < 0.01, ***p < 0.001, ****p < 0.0001. Notable near-significant differences (0.05 < p < 0.1) are indicated in the figures. Notable non-significant (and non-near significant) differences are indicated in the figures by ‘‘n.s”.

### Data availability

The data that support the findings of this study are available from the corresponding author upon request.

## REFERENCES

1. Kupfer, D. J., Frank, E. & Phillips, M. L. Major depressive disorder: new clinical, neurobiological, and treatment perspectives. Lancet 379, 1045–1055 (2012).

2. Sheline, Y. I., Wang, P. W., Gado, M. H., Csernansky, J. G. & Vannier, M. W. Hippocampal atrophy in recurrent major depression. Proc. Natl. Acad. Sci. U. S. A. (1996).

3. Campbell, S. & MacQueen, G. An update on regional brain volume differences associated with mood disorders. Current Opinion in Psychiatry (2006). doi:10.1097/01.yco.0000194371.47685.f2

4. Shen, Z. et al. Changes of grey matter volume in first-episode drug-naive adult major depressive disorder patients with different age-onset. NeuroImage Clin. (2016). doi:10.1016/j.nicl.2016.08.016

5. Cameron, H. A. & Gould, E. Adult neurogenesis is regulated by adrenal steroids in the dentate gyrus. Neuroscience (1994). doi:10.1016/0306-4522(94)90224-0

6. Sahay, A. & Hen, R. Adult hippocampal neurogenesis in depression. Nat. Neurosci. (2007). doi:10.1038/nn1969

7. Schoenfeld, T. J. & Gould, E. Stress, stress hormones, and adult neurogenesis. Experimental Neurology (2012). doi:10.1016/j.expneurol.2011.01.008

8. Sheline, Y. I., Liston, C. & McEwen, B. S. Parsing the Hippocampus in Depression: Chronic Stress, Hippocampal Volume, and Major Depressive Disorder. Biol. Psychiatry (2019). doi:10.1016/j.biopsych.2019.01.011

9. Snyder, J. S., Soumier, A., Brewer, M., Pickel, J. & Cameron, H. A. Adult hippocampal neurogenesis buffers stress responses and depressive behaviour. Nature 476, 458–61 (2011).

10. Glover, L. R., Schoenfeld, T. J., Karlsson, R. M., Bannerman, D. M. & Cameron, H. A. Ongoing neurogenesis in the adult dentate gyrus mediates behavioral responses to ambiguous threat cues. PLoS Biol. (2017). doi:10.1371/journal.pbio.2001154

11. Egeland, M., Zunszain, P. A. & Pariante, C. M. Molecular mechanisms in the regulation of adult neurogenesis during stress. Nature Reviews Neuroscience (2015). doi:10.1038/nrn3855

12. Santarelli, L. et al. Requirement of Hippocampal Neurogenesis for the Behavioral Effects of Antidepressants. Science (80-.). 301, 805–809 (2003).

13. Surget, A. et al. Antidepressants recruit new neurons to improve stress response regulation. Mol. Psychiatry (2011). doi:10.1038/mp.2011.48

14. Culig, L. et al. Increasing adult hippocampal neurogenesis in mice after exposure to unpredictable chronic mild stress may counteract some of the effects of stress. Neuropharmacology (2017). doi:10.1016/j.neuropharm.2017.09.009

15. Jacobs, B. L., Van Praag, H. & Gage, F. H. Adult brain neurogenesis and psychiatry: A novel theory of depression. Molecular Psychiatry (2000). doi:10.1038/sj.mp.4000712

16. Miller, B. R. & Hen, R. The current state of the neurogenic theory of depression and anxiety. Current Opinion in Neurobiology (2015). doi:10.1016/j.conb.2014.08.012

17. Belkaid, Y. & Hand, T. W. Role of the microbiota in immunity and inflammation. Cell 157, 121–41 (2014).

18. Cani, P. D. Metabolism in 2013: The gut microbiota manages host metabolism. Nat. Rev. Endocrinol. 10, 74–6 (2014).

19. Sharon, G. et al. The Central Nervous System and the Gut Microbiome. Cell 167, 915–932 (2016).

20. Jiang, H. et al. Altered fecal microbiota composition in patients with major depressive disorder. Brain. Behav. Immun. 48, 186–94 (2015).

21. Naseribafrouei, A. et al. Correlation between the human fecal microbiota and depression. Neurogastroenterol. Motil. 26, 1155–1162 (2014).

22. Bercik, P. et al. The intestinal microbiota affect central levels of brain-derived neurotropic factor and behavior in mice. Gastroenterology 141, 599–609, 609.e1–3 (2011).

23. Bravo, J. A. et al. Ingestion of Lactobacillus strain regulates emotional behavior and central GABA receptor expression in a mouse via the vagus nerve. Proc. Natl. Acad. Sci. U. S. A. 108, 16050–5 (2011).

24. Heijtz, R. D. et al. Normal gut microbiota modulates brain development and behavior. Proc. Natl. Acad. Sci. U. S. A. 108, 3047–52 (2011).

25. Hsiao, E. Y. et al. Microbiota modulate behavioral and physiological abnormalities associated with neurodevelopmental disorders. Cell 155, 1451–63 (2013).

26. Sampson, T. R. et al. Gut Microbiota Regulate Motor Deficits and Neuroinflammation in a Model of Parkinson’s Disease. Cell 167, 1469–1480.e12 (2016).

27. Schroeder, B. O. & Bäckhed, F. Signals from the gut microbiota to distant organs in physiology and disease. Nat. Med. 22, 1079–1089 (2016).

28. Kelly, J. R. et al. Transferring the blues: Depression-associated gut microbiota induces neurobehavioural changes in the rat. J. Psychiatr. Res. 82, 109–18 (2016).

29. Hodes, G. E., Kana, V., Menard, C., Merad, M. & Russo, S. J. Neuroimmune mechanisms of depression. Nat. Neurosci. 18, 1386–93 (2015).

30. Parker, K. J., Schatzberg, A. F. & Lyons, D. M. Neuroendocrine aspects of hypercortisolism in major depression. Horm. Behav. 43, 60–6 (2003).

31. McEwen, B. S. Physiology and neurobiology of stress and adaptation: central role of the brain. Physiol. Rev. 87, 873–904 (2007).

32. Sahay, A. & Hen, R. Adult hippocampal neurogenesis in depression. Nat. Neurosci. 10, 1110–1115 (2007).

33. Kennedy, P. J. et al. Gut memories: towards a cognitive neurobiology of irritable bowel syndrome. Neurosci. Biobehav. Rev. 36, 310–40 (2012).

34. Burokas, A. et al. Targeting the Microbiota-Gut-Brain Axis: Prebiotics Have Anxiolytic and Antidepressant-like Effects and Reverse the Impact of Chronic Stress in Mice. Biol. Psychiatry 82, 472–487 (2017).

35. Savignac, H. M., Kiely, B., Dinan, T. G. & Cryan, J. F. Bifidobacteria exert strain-specific effects on stress-related behavior and physiology in BALB/c mice. Neurogastroenterol. Motil. 26, 1615–27 (2014).

36. Yang, C. et al. Bifidobacterium in the gut microbiota confer resilience to chronic social defeat stress in mice. Sci. Rep. 7, 45942 (2017).

37. Pirbaglou, M. et al. Probiotic supplementation can positively affect anxiety and depressive symptoms: a systematic review of randomized controlled trials. Nutrition Research (2016). doi:10.1016/j.nutres.2016.06.009

38. Huang, R., Wang, K. & Hu, J. Effect of probiotics on depression: A systematic review and meta-analysis of randomized controlled trials. Nutrients (2016). doi:10.3390/nu8080483

39. Ng, Q. X., Peters, C., Ho, C. Y. X., Lim, D. Y. & Yeo, W.-S. A meta-analysis of the use of probiotics to alleviate depressive symptoms. J. Affect. Disord. (2018). doi:10.1016/j.jad.2017.11.063

40. Willner, P. Validity, reliability and utility of the chronic mild stress model of depression: a 10-year review and evaluation. Psychopharmacology (Berl). 134, 319–29 (1997).

41. Willner, P. Chronic mild stress (CMS) revisited: consistency and behavioural-neurobiological concordance in the effects of CMS. Neuropsychobiology 52, 90–110 (2005).

42. Nollet, M., Guisquet, A.-M. Le & Belzung, C. Models of Depression: Unpredictable Chronic Mild Stress in Mice. Curr. Protoc. Pharmacol. 61, 5.65.1–5.65.17 (2013).

43. Monteiro, S. et al. An efficient chronic unpredictable stress protocol to induce stress-related responses in C57BL/6 mice. Front. psychiatry 6, 6 (2015).

44. Hill, M. N. et al. The therapeutic potential of the endocannabinoid system for the development of a novel class of antidepressants. Trends Pharmacol. Sci. 30, 484–93 (2009).

45. Lutz, B. Endocannabinoid signals in the control of emotion. Curr. Opin. Pharmacol. 9, 46–52 (2009).

46. Aguado, T. et al. The endocannabinoid system drives neural progenitor proliferation. FASEB J. 19, 1704–6 (2005).

47. Aguado, T. et al. The CB1 cannabinoid receptor mediates excitotoxicity-induced neural progenitor proliferation and neurogenesis. J. Biol. Chem. 282, 23892–8 (2007).

48. Jin, K. et al. Defective adult neurogenesis in CB1 cannabinoid receptor knockout mice. Mol. Pharmacol. 66, 204–8 (2004).

49. Dócs, K. et al. The Ratio of 2-AG to Its Isomer 1-AG as an Intrinsic Fine Tuning Mechanism of CB1 Receptor Activation. Front. Cell. Neurosci. (2017). doi:10.3389/fncel.2017.00039

50. Jefferies, H. B. et al. Rapamycin suppresses 5’TOP mRNA translation through inhibition of p70s6k. EMBO J. 16, 3693–704 (1997).

51. Proud, C. G. Signalling to translation: how signal transduction pathways control the protein synthetic machinery. Biochem. J. 403, 217–34 (2007).

52. Long, J. Z. et al. Selective blockade of 2-arachidonoylglycerol hydrolysis produces cannabinoid behavioral effects. Nat. Chem. Biol. 5, 37–44 (2009).

53. Pan, B. et al. Blockade of 2-Arachidonoylglycerol Hydrolysis by Selective Monoacylglycerol Lipase Inhibitor 4-Nitrophenyl 4-(Dibenzo[d][1,3]dioxol-5-yl(hydroxy)methyl)piperidine-1-carboxylate (JZL184) Enhances Retrograde Endocannabinoid Signaling. J. Pharmacol. Exp. Ther. 331, 591–597 (2009).

54. Tam, J. et al. Peripheral CB1 cannabinoid receptor blockade improves cardiometabolic risk in mouse models of obesity. J. Clin. Invest. 120, 2953–66 (2010).

55. Anacker, C. et al. Hippocampal neurogenesis confers stress resilience by inhibiting the ventral dentate gyrus. Nature (2018). doi:10.1038/s41586-018-0262-4

56. Galley, J. D. et al. Exposure to a social stressor disrupts the community structure of the colonic mucosa-associated microbiota. BMC Microbiol. 14, 189 (2014).

57. Jašarević, E., Rodgers, A. B. & Bale, T. L. A novel role for maternal stress and microbial transmission in early life programming and neurodevelopment. Neurobiol. Stress 1, 81–88 (2015).

58. Marin, I. A. et al. Microbiota alteration is associated with the development of stress-induced despair behavior. Sci. Rep. 7, 43859 (2017).

59. Aizawa, E. et al. Possible association of Bifidobacterium and Lactobacillus in the gut microbiota of patients with major depressive disorder. J. Affect. Disord. 202, 254–7 (2016).

60. Salaj, R. et al. The effects of two Lactobacillus plantarum strains on rat lipid metabolism receiving a high fat diet. ScientificWorldJournal. 2013, 135142 (2013).

61. Wu, Y., Zhang, Q., Ren, Y. & Ruan, Z. Effect of probiotic Lactobacillus on lipid profile: A systematic review and meta-analysis of randomized, controlled trials. PLoS One 12, e0178868 (2017).

62. Schwarzer, M. et al. Lactobacillus plantarum strain maintains growth of infant mice during chronic undernutrition. Science (80-.). 351, 854–857 (2016).

63. Liu, Y.-W. et al. Psychotropic effects of Lactobacillus plantarum PS128 in early life-stressed and naïve adult mice. Brain Res. 1631, 1–12 (2016).

64. Mazure, C. M. Life Stressors as Risk Factors in Depression. Clin. Psychol. Sci. Pract. 5, 291–313 (1998).

65. Overstreet, D. H. Modeling depression in animal models. Methods Mol. Biol. 829, 125–44 (2012).

66. Moreira, F. A. & Crippa, J. A. S. The psychiatric side-effects of rimonabant. Rev. Bras. Psiquiatr. 31, 145–53 (2009).

67. Monteleone, P. et al. Investigation of CNR1 and FAAH endocannabinoid gene polymorphisms in bipolar disorder and major depression. Pharmacol. Res. 61, 400–4 (2010).

68. Denson, T. F. & Earleywine, M. Decreased depression in marijuana users. Addict. Behav. 31, 738–742 (2006).

69. Jiang, W. et al. Cannabinoids promote embryonic and adult hippocampus neurogenesis and produce anxiolytic- and antidepressant-like effects. J. Clin. Invest. 115, 3104–16 (2005).

70. Zhong, P. et al. Monoacylglycerol lipase inhibition blocks chronic stress-induced depressive-like behaviors via activation of mTOR signaling. Neuropsychopharmacology 39, 1763–76 (2014).

71. Hill, M. N. et al. Downregulation of endocannabinoid signaling in the hippocampus following chronic unpredictable stress. Neuropsychopharmacology 30, 508–15 (2005).

72. Wang, W. et al. Deficiency in endocannabinoid signaling in the nucleus accumbens induced by chronic unpredictable stress. Neuropsychopharmacology 35, 2249–61 (2010).

73. Cravatt, B. F. et al. Molecular characterization of an enzyme that degrades neuromodulatory fatty-acid amides. Nature 384, 83–7 (1996).

74. Blankman, J. L., Simon, G. M. & Cravatt, B. F. A comprehensive profile of brain enzymes that hydrolyze the endocannabinoid 2-arachidonoylglycerol. Chem. Biol. 14, 1347–56 (2007).

75. McLaughlin, R. J., Hill, M. N., Morrish, A. C. & Gorzalka, B. B. Local enhancement of cannabinoid CB1 receptor signalling in the dorsal hippocampus elicits an antidepressant-like effect. Behav. Pharmacol. 18, 431– 8 (2007).

76. Duric, V. et al. A negative regulator of MAP kinase causes depressive behavior. Nat. Med. 16, 1328–32 (2010).

77. Jernigan, C. S. et al. The mTOR signaling pathway in the prefrontal cortex is compromised in major depressive disorder. Prog. Neuropsychopharmacol. Biol. Psychiatry 35, 1774–9 (2011).

78. Hill, M. N. & Gorzalka, B. B. Impairments in Endocannabinoid Signaling and Depressive Illness. JAMA 301, 1165 (2009).

79. Hill, M. N., Miller, G. E., Ho, W.-S. V, Gorzalka, B. B. & Hillard, C. J. Serum endocannabinoid content is altered in females with depressive disorders: a preliminary report. Pharmacopsychiatry 41, 48–53 (2008).

80. Yi, B. et al. Reductions in circulating endocannabinoid 2-arachidonoylglycerol levels in healthy human subjects exposed to chronic stressors. Prog. Neuropsychopharmacol. Biol. Psychiatry 67, 92–7 (2016).

81. Rueda, D., Navarro, B., Martínez-Serrano, A., Guzmán, M. & Galve-Roperh, I. The Endocannabinoid Anandamide Inhibits Neuronal Progenitor Cell Differentiation through Attenuation of the Rap1/B-Raf/ERK Pathway. J. Biol. Chem. 277, 46645–46650 (2002).

82. Puighermanal, E. et al. Cannabinoid modulation of hippocampal long-term memory is mediated by mTOR signaling. Nat. Neurosci. 12, 1152–8 (2009).

83. Prenderville, J. A., Kelly, Á. M. & Downer, E. J. The role of cannabinoids in adult neurogenesis. Br. J. Pharmacol. 172, 3950–3963 (2015).

84. Cani, P. D. et al. Endocannabinoids - at the crossroads between the gut microbiota and host metabolism. Nat. Rev. Endocrinol. 12, 133–43 (2016).

85. Muccioli, G. G. et al. The endocannabinoid system links gut microbiota to adipogenesis. Mol. Syst. Biol. 6, 392 (2010).

86. Everard, A. et al. Cross-talk between Akkermansia muciniphila and intestinal epithelium controls diet-induced obesity. Proc. Natl. Acad. Sci. (2013). doi:10.1073/pnas.1219451110

87. Guida, F. et al. Antibiotic-induced microbiota perturbation causes gut endocannabinoidome changes, hippocampal neuroglial reorganization and depression in mice. Brain. Behav. Immun. (2018). doi:10.1016/j.bbi.2017.09.001

88. Bailey, M. T. et al. Exposure to a social stressor alters the structure of the intestinal microbiota: implications for stressor-induced immunomodulation. Brain. Behav. Immun. 25, 397–407 (2011).

89. Jašarević, E., Howerton, C. L., Howard, C. D. & Bale, T. L. Alterations in the Vaginal Microbiome by Maternal Stress Are Associated With Metabolic Reprogramming of the Offspring Gut and Brain. Endocrinology 156, 3265– 3276 (2015).

90. De Palma, G. et al. Microbiota and host determinants of behavioural phenotype in maternally separated mice. Nat. Commun. 6, 7735 (2015).

91. Zheng, P. et al. Gut microbiome remodeling induces depressive-like behaviors through a pathway mediated by the host’s metabolism. Mol. Psychiatry 21, 786–796 (2016).

92. Möhle, L. et al. Ly6Chi Monocytes Provide a Link between Antibiotic-Induced Changes in Gut Microbiota and Adult Hippocampal Neurogenesis. Cell Rep. 15, 1945–1956 (2016).

93. Sawada, N. et al. Regulation by commensal bacteria of neurogenesis in the subventricular zone of adult mouse brain. Biochem. Biophys. Res. Commun. 498, 824–829 (2018).

94. Dinan, T. G. & Cryan, J. F. Melancholic microbes: a link between gut microbiota and depression? Neurogastroenterol. Motil. 25, 713–9 (2013).

95. Foster, J. A. & McVey Neufeld, K.-A. Gut-brain axis: how the microbiome influences anxiety and depression. Trends Neurosci. 36, 305–12 (2013).

96. Sarkar, A. et al. Psychobiotics and the Manipulation of Bacteria–Gut–Brain Signals. Trends Neurosci. 39, 763–781 (2016).

97. Rao, A. V. et al. A randomized, double-blind, placebo-controlled pilot study of a probiotic in emotional symptoms of chronic fatigue syndrome. Gut Pathog. 1, 6 (2009).

98. Slykerman, R. F. et al. Effect of Lactobacillus rhamnosus HN001 in Pregnancy on Postpartum Symptoms of Depression and Anxiety: A Randomised Double-blind Placebo-controlled Trial. EBioMedicine 24, 159–165 (2017).

99. Akkasheh, G. et al. Clinical and metabolic response to probiotic administration in patients with major depressive disorder: A randomized, double-blind, placebo-controlled trial. Nutrition 32, 315–20 (2016).

100. Bambling, M., Edwards, S. C., Hall, S. & Vitetta, L. A combination of probiotics and magnesium orotate attenuate depression in a small SSRI resistant cohort: an intestinal anti-inflammatory response is suggested. Inflammopharmacology 25, 271–274 (2017).

101. Lew, L.-C. et al. Probiotic Lactobacillus plantarum P8 alleviated stress and anxiety while enhancing memory and cognition in stressed adults: A randomised, double-blind, placebo-controlled study. Clin. Nutr. (2018). doi:10.1016/J.CLNU.2018.09.010

102. Lafourcade, M. et al. Nutritional omega-3 deficiency abolishes endocannabinoid-mediated neuronal functions. Nat. Neurosci. 14, 345–50 (2011).

103. Vancassel, S. et al. n-3 polyunsaturated fatty acid supplementation reverses stress-induced modifications on brain monoamine levels in mice. J. Lipid Res. 49, 340–8 (2008).

104. Falcinelli, S. et al. Lactobacillus rhamnosus lowers zebrafish lipid content by changing gut microbiota and host transcription of genes involved in lipid metabolism. Sci. Rep. 5, 9336 (2015).

105. Semova, I. et al. Microbiota regulate intestinal absorption and metabolism of fatty acids in the zebrafish. Cell Host Microbe 12, 277–88 (2012).

106. Chiu, C.-H., Lu, T.-Y., Tseng, Y.-Y. & Pan, T.-M. The effects of Lactobacillus-fermented milk on lipid metabolism in hamsters fed on high-cholesterol diet. Appl. Microbiol. Biotechnol. 71, 238–45 (2006).

107. Kishino, S. et al. Polyunsaturated fatty acid saturation by gut lactic acid bacteria affecting host lipid composition. Proc. Natl. Acad. Sci. U. S. A. 110, 17808–13 (2013).

108. Bao, Y. et al. Effect of *Lactobacillus plantarum* P-8 on lipid metabolism in hyperlipidemic rat model. Eur. J. Lipid Sci. Technol. 114, 1230–1236 (2012).

109. Xie, N. et al. Effects of two Lactobacillus strains on lipid metabolism and intestinal microflora in rats fed a high-cholesterol diet. BMC Complement. Altern. Med. 11, 53 (2011).

## REFERENCES for METHODS

1. Long, J. Z. et al. Selective blockade of 2-arachidonoylglycerol hydrolysis produces cannabinoid behavioral effects. Nat. Chem. Biol. 5, 37–44 (2009).

2. Zhong, P. et al. Monoacylglycerol lipase inhibition blocks chronic stress-induced depressive-like behaviors via activation of mTOR signaling. Neuropsychopharmacology 39, 1763–76 (2014).

3. Kinsey, S. G. et al. Repeated low-dose administration of the monoacylglycerol lipase inhibitor JZL184 retains cannabinoid receptor type 1-mediated antinociceptive and gastroprotective effects. J. Pharmacol. Exp. Ther. 345, 492– 501 (2013).

4. Quereda, J. J. et al. Bacteriocin from epidemic Listeria strains alters the host intestinal microbiota to favor infection. Proc. Natl. Acad. Sci. U. S. A. 113, 5706– 11 (2016).

5. Cole, J. R. et al. The Ribosomal Database Project: improved alignments and new tools for rRNA analysis. Nucleic Acids Res. 37, D141–5 (2009).

6. Dickson, L. B. et al. Carryover effects of larval exposure to different environmental bacteria drive adult trait variation in a mosquito vector. Sci. Adv. 3, e1700585 (2017).

7. Benjamini, Y. & Hochberg, Y. Controlling the False Discovery Rate: A Practical and Powerful Approach to Multiple Testing. Journal of the Royal Statistical Society. Series B (Methodological) 57, 289–300 (1995).

8. Ducottet, C., Aubert, A. & Belzung, C. Susceptibility to subchronic unpredictable stress is related to individual reactivity to threat stimuli in mice. Behav. Brain Res. 155, 291–9 (2004).

9. Nollet, M., Guisquet, A.-M. Le & Belzung, C. Models of Depression: Unpredictable Chronic Mild Stress in Mice. Curr. Protoc. Pharmacol. 61, 5.65.1–5.65.17 (2013).

10. Santarelli, L. et al. Requirement of Hippocampal Neurogenesis for the Behavioral Effects of Antidepressants. Science (80). 301, 805–809 (2003).

11. de Chaumont, F. et al. Icy: an open bioimage informatics platform for extended reproducible research. Nat. Methods 9, 690–6 (2012).

12. Wang, W. et al. Deficiency in endocannabinoid signaling in the nucleus accumbens induced by chronic unpredictable stress. Neuropsychopharmacology 35, 2249–61 (2010).

13. Patel, S., Rademacher, D. J. & Hillard, C. J. Differential regulation of the endocannabinoids anandamide and 2-arachidonylglycerol within the limbic forebrain by dopamine receptor activity. J. Pharmacol. Exp. Ther. 306, 880–8 (2003).

