## Supplementary figures for "The impact of gut microbiota on depressive-like behaviors and adult hippocampal neurogenesis requires the endocannabinoid system"

**Supplementary Materials (5 figures and 1 table)**

Supplementary Table 1 : Timeline of the stressor exposure used in the Unpredictable Chronic Mild Stress (UCMS) protocol.

Supplementary Figure 1: Microbiota from UCMS Mice Transfers Depressive-like behaviors.

Supplementary Figure 2 : UCMS Microbiota Transfers Depressive-like Behaviors to Recipient Antibiotic-treated Mice.

Supplementary Figure 3 : Complementary Analysis of Metabolomic Profiles in the Serum of Recipient Mice.

Supplementary Figure 4 : Adult neurogenesis in dorsal and ventral hippocampus are modulated by the eCB system.

Supplementary Figure 5 : Unpredictable Chronic Mild Stress (UCMS) Alters Gut Microbiota and is Transferable to Recipient Mice.

Supplementary Table 1

| | Monday | Tuesday | Wednesday | Thursday | Friday |
|---|---|---|---|---|---|
| Week 1 | Blood & feces (11h-13h) | No sawdust<br>(10h-14h) | 3 cage changes<br>(9h30-12h30) | Damp sawdust<br>(10h-13h30h) | No sawdust (10h-13h) |
| | Cage tilt 45° (14h30-20h) | Damp sawdust (14h-18h) | Foreign odor (18h) | Cage tilt (14h-18h) | Foreign odor (14h)<br>Reversal of the<br>L/D cycle (19h) |
| Week 2 | Back to normal L/D cycle<br>(9h) | Damp sawdust<br>(10h-16h) | No sawdust<br>(10h30-15h30) | Cage tilt at 45°<br>(10h) | Damp sawdust (10h-13h) |
| | Confinement (14h-15h)<br>Foreign odor (15h) | Cage tilt at 45°<br>(18h) | Foreign odor<br>(15h30) | Confinement<br>(15h30-16h30)<br>Damp sawdust (16h30) | Foreign cage (14h)<br>Reversal of the<br>L/D cycle (19h) |
| Week 3 | Back to normal L/D cycle<br>Cage tilt at 45°(10h) | No sawdust (10h-14h) | No sawdust + Water<br>(10h-15h30) | Confinement (10h30-12h) | Damp sawdust (10h-13h) |
| | Damp sawdust (14h-18h) | Confinement (14h-15h30) | Cage tilt (14h) | 4 cage changes (14h-18h)<br>Foreign odor (18h) | No sawdust (13h-19h)<br>Reversal of the<br>L/D cycle (19h) |
| Week 4 | Back to normal L/D cycle<br>(9h30)<br>Cage tilt at 45° (9h30-14h) | No sawdust<br>(10h-14h) | 3 cage changes<br>(9h30-12h30) | Damp sawdust<br>(10h-13h30h) | No sawdust (10h-14h) |
| | Confinement (14h-15h30) | Damp sawdust<br>(14h-18h) | Cage tilt 45° (12h30-18h)<br>Foreign odor (18h) | Confinement<br>(13h30-15h00) | Cage tilt 45° (14h)<br>Reversal of the<br>L/D cycle (19h) |

Repeated once for a total of 8 weeks

Supplementary Figure 1

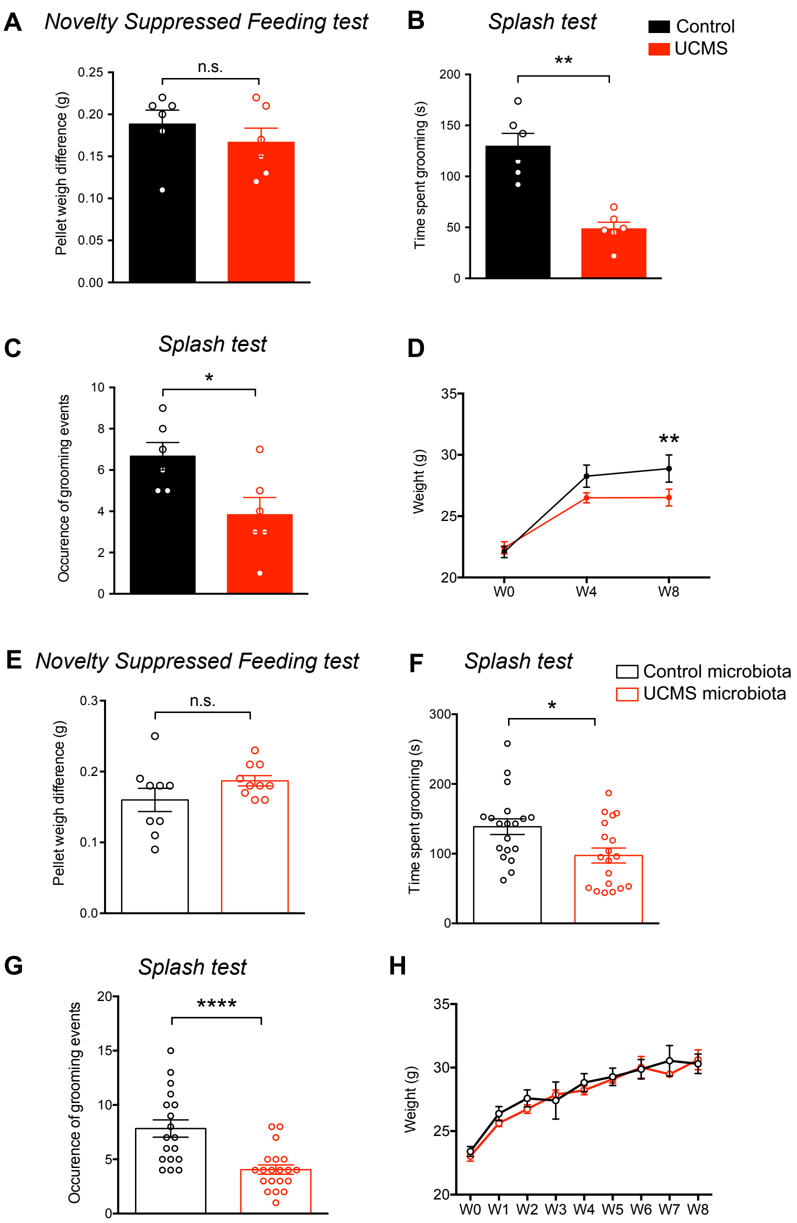


**Supplementary Figure 1. Microbiota from UCMS Mice Transfers Depressive-like Behaviors**. Control mice, or mice subjected to UCMS (donor mice) (**A-D**), and mice recipient of the microbiota from control or UCMS mice (**E-H**), underwent different behavioral tests. **A**,**E**, Feeding drive of Control mice ($n$ = 6), UCMS mice ($n$ = 6), Control microbiota- ($n$ = 9) and UCMS microbiota-recipient mice ($n$ = 10) in the novelty suppressed feeding test as assessed by the pellet consumption (weight difference)

during 10 minutes after the test (Control vs UCMS mice, *P* = 0.5152; Control microbiota- vs UCMS microbiota-recipient mice, *P* = 0.2271). **B,F,** Time spent grooming for Control mice (*n* = 6), UCMS mice (*n* = 6), Control microbiota- (*n* = 19) and UCMS microbiota-recipient mice (*n* = 19) in the splash test (Control vs UCMS mice, *P* = 0.0022; Control microbiota- vs UCMS microbiota-recipient mice, *P* = 0.0210). **C,G**, Occurrence of self-grooming events for Control mice (*n* = 6), UCMS mice (*n* = 6), Control microbiota- (*n* = 19) and UCMS microbiota-recipient mice (*n* = 19) in the splash test (Control vs UCMS mice, *P* = 0.0325; Control microbiota- vs UCMS microbiota-recipient mice, *P* < 0.0001). **D,H**, Body weight was measured during the UCMS protocol (*n* = 6/group, **D**) and after FMT in Control microbiota (*n* = 9) and UCMS microbiota-recipient mice (*n* = 10, **H**). Data are represented as mean ± s.e.m. Statistical significance was calculated using the Mann Whitney test (**P* < 0.05, ** *P* < 0.01, *** *P* < 0.001, **** *P* < 0.0001).

Supplementary Figure 2

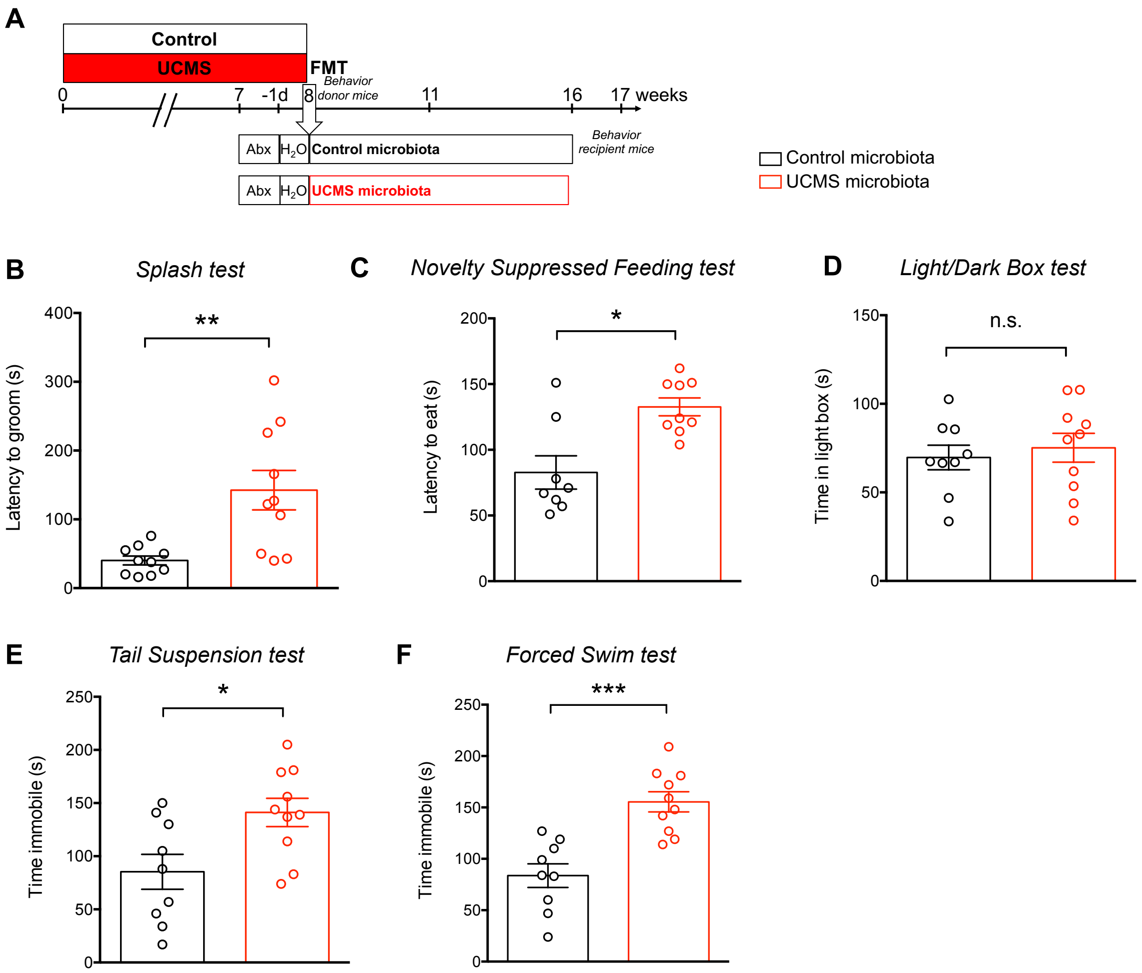


**Supplementary Figure 2. Unpredictable Chronic Mild Stress (UCMS) Microbiota Transfers Depressive-like Behaviors to Recipient Antibiotic-treated Mice. A,** Experimental timeline of Fecal Microbiota Transplantation (FMT) from Control and UCMS mice, respectively ‘Control microbiota' and ‘UCMS microbiota’, to antibiotic (Abx)-treated recipient mice. Recipient mice were given a combination of vancomycin (0.5 g/l), ampicillin (1 g/l), streptomycin (5 g/L), colistin (1 g/l), and metronidazole (0.5 g/l) in their drinking water for 6 consecutive days. Twenty-four hours later, animals were colonized via two rounds of oral gavage with microbiota, separated 3 days apart, and kept in separate sterile isolators. Donor microbiota was acquired from pooled fecal samples from 5-6 animals and resuspended in PBS. **B-F,** Recipient mice underwent different behavioral tests: **B,** Latency to groom in the splash test for Control microbiota- (*n* = 10) and UCMS microbiota-recipient mice (*n* = 10) (Control microbiota- vs UCMS microbiota-recipient mice, $P = 0.0026$). **C,** Latency to eat in a novel environment in the Novelty Suppressed Feeding test for Control microbiota- (*n* = 8), UCMS microbiota-recipient mice (*n* = 10) (Control microbiota- vs UCMS microbiota-recipient mice, $P = 0.0214$). **D,** Time spent in the light box in the Light/Dark Box test for Control microbiota- (*n* = 9) and UCMS microbiota-recipient mice (*n* = 10) (Control microbiota- vs UCMS microbiota-recipient mice, $P = 0.6038$). **E,F**, Time spent immobile in the Tail Suspension test (**E**) and time spent immobile in the Forced Swim Test for Control microbiota- (*n* = 9,) and UCMS microbiota-recipient mice (*n* = 10, **F**) (Control microbiota- vs UCMS microbiota-recipient mice, $P = 0.0002$). Data are represented as mean ± s.e.m. Statistical significance was calculated using the Mann Whitney test (*$P < 0.05$, ** $P < 0.01$, *** $P < 0.001$).

Supplementary Figure 3

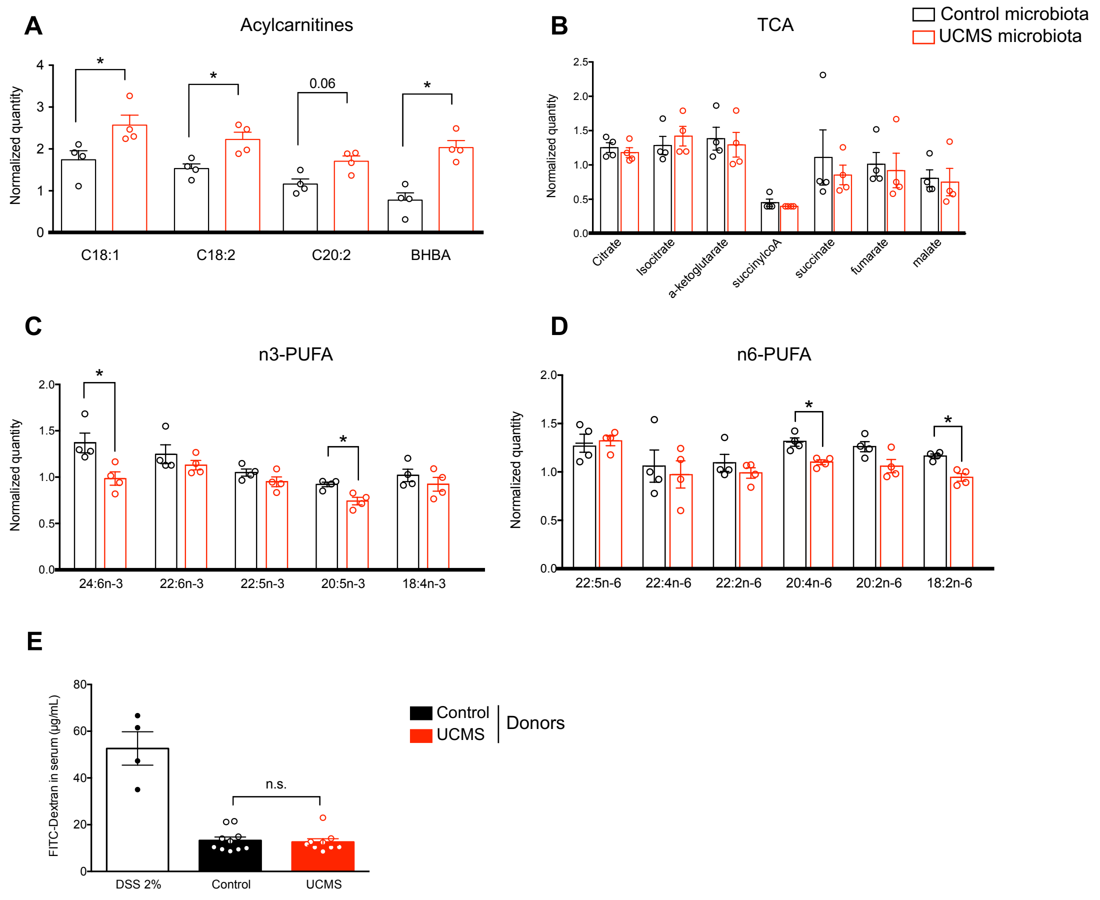


**Supplementary Figure 3. Complementary analysis of metabolomic profiles in the Serum of Recipient Mice. A-D.** The serum was analyzed by mass spectrometry for small molecules. Shown are normalized levels of acylcarnitines (*, $P = 0.0286$; except for C20:2 with $P = 0.0571$, **A**), products of the TCA cycle (**B**), n3-PUFA (*, $P = 0.0286$) (**C**) and n6-PUFA (*, $P = 0.0286$) (**D**) ($n = 4$/group). Data are represented as mean ± s.e.m. Statistical significance was calculated using the Mann Whitney test (* $P < 0.05$). **E,** Intestinal permeability was measured by FITC intensity in serum after oral gavage with FITC-Dextran in Control ($n = 10$) and UCMS mice ($n = 9$). Dextran sodium sulfate (DSS) is used as a positive control ($n = 4$). Data are represented as mean ± s.e.m. Mann Whitney test (Control vs UCMS mice, $P > 0.999$).

Supplementary Figure 4

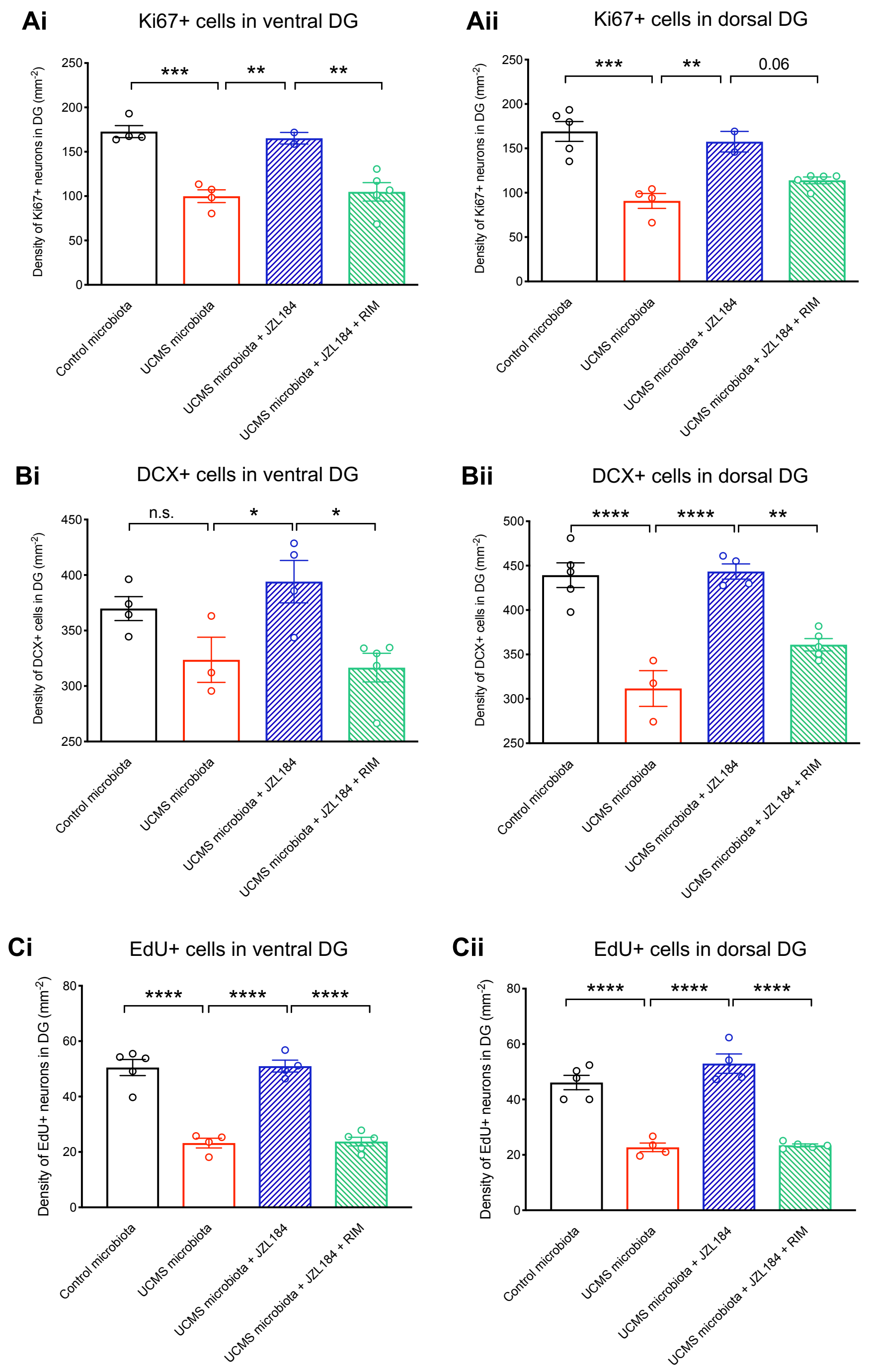


**Supplementary Figure 4. Adult neurogenesis in dorsal and ventral hippocampus are modulated by the eCB system.**

**Ai**, Quantitative evaluation of the density of Ki67$^{+}$ cells in ventral DG for Control microbiota-recipient mice ($n$ = 4), UCMS microbiota-recipient mice ($n$ = 4), UCMS microbiota-recipient mice treated with JZL184 (*n* = 2) and UCMS microbiota-recipient mice treated with JZL184 and rimonabant (*n* = 5). (Control microbiota- vs UCMS microbiota-recipient mice, *P* = 0.0005; UCMS microbiota-recipient mice vs UCMS microbiota-recipient mice + JZL184, *P* = 0.006; UCMS microbiota-recipient mice + JZL184 vs UCMS microbiota-recipient mice + JZL184 + RIM, *P* = 0.0081). **Aii**, Quantitative evaluation of the density of Ki67$^+$ cells in dorsal DG for Control microbiota-recipient mice (*n* = 5), UCMS microbiota-recipient mice (*n* = 4), UCMS microbiota-recipient mice treated with JZL184 (*n* = 2), UCMS microbiota-recipient mice treated with JZL184 and rimonabant (*n* = 5). (Control microbiota- vs UCMS microbiota-recipient mice, *P* = 0.0002; UCMS microbiota-recipient mice vs UCMS microbiota-recipient mice + JZL184, *P* = 0.0053; UCMS microbiota-recipient mice + JZL184 vs UCMS microbiota-recipient mice + JZL184 + RIM, *P* = 0.0586). **Bi**, Quantitative evaluation of the density of DCX$^+$ cells in ventral DG for Control microbiota-recipient mice (*n* = 4), UCMS microbiota-recipient mice (*n* = 3), UCMS microbiota-recipient mice treated with JZL184 (*n* = 4), UCMS microbiota-recipient mice treated with JZL184 and rimonabant (*n* = 5). Control microbiota- vs UCMS microbiota-recipient mice, *P* = 0.2603; UCMS microbiota-recipient mice vs UCMS microbiota-recipient mice + JZL184, *P* = 0.0498; UCMS microbiota-recipient mice + JZL184 vs UCMS microbiota-recipient mice + JZL184 + RIM, *P* = 0.0135. **Bii**, Quantitative evaluation of the density of DCX$^+$ cells in dorsal DG for Control microbiota-recipient mice (*n* = 5), UCMS microbiota-recipient mice (*n* = 3), UCMS microbiota-recipient mice treated with JZL184 (*n* = 4) and UCMS microbiota-recipient mice treated with JZL184 and rimonabant (*n* = 5). Control microbiota- vs UCMS microbiota-recipient mice, *P* < 0.0001; UCMS microbiota-recipient mice vs UCMS microbiota-recipient mice + JZL184, *P* < 0.0001; UCMS microbiota-recipient mice + JZL184 vs UCMS microbiota-recipient mice + JZL184 + RIM, *P* = 0.0014. **Ci**,

Quantitative evaluation of the density of $EdU^+$ cells in ventral DG for Control microbiota-recipient mice ($n = 5$), UCMS microbiota-recipient mice ($n = 4$), UCMS microbiota-recipient mice treated with JZL184 ($n = 4$), UCMS microbiota-recipient mice treated with JZL184 and rimonabant ($n = 5$). Control microbiota- vs UCMS microbiota-recipient mice, $P < 0.0001$; UCMS microbiota-recipient mice vs UCMS microbiota-recipient mice + JZL184, $P < 0.0001$; UCMS microbiota-recipient mice + JZL184 vs UCMS microbiota-recipient mice + JZL184 + RIM, $P < 0.0001$. **Cii,** Quantitative evaluation of the density of $EdU^+$ cells in dorsal DG for Control microbiota-recipient mice ($n = 5$), UCMS microbiota-recipient mice ($n = 4$), UCMS microbiota-recipient mice treated with JZL184 ($n = 4$) and UCMS microbiota-recipient mice treated with JZL184 and rimonabant ($n = 5$). Control microbiota- vs UCMS microbiota-recipient mice, $P < 0.0001$; UCMS microbiota-recipient mice vs UCMS microbiota-recipient mice + JZL184, $P < 0.0001$; UCMS microbiota-recipient mice + JZL184 vs UCMS microbiota-recipient mice + JZL184 + RIM, $P < 0.0001$. Scale bars: 100µm. Data are represented as mean ± s.e.m. Statistical significance was calculated using One-way ANOVA with Tukey's multiple comparisons test (*$P < 0.05$, ** $P < 0.01$, *** $P < 0.005$, **** $P < 0.0001$).

Supplementary Figure 5

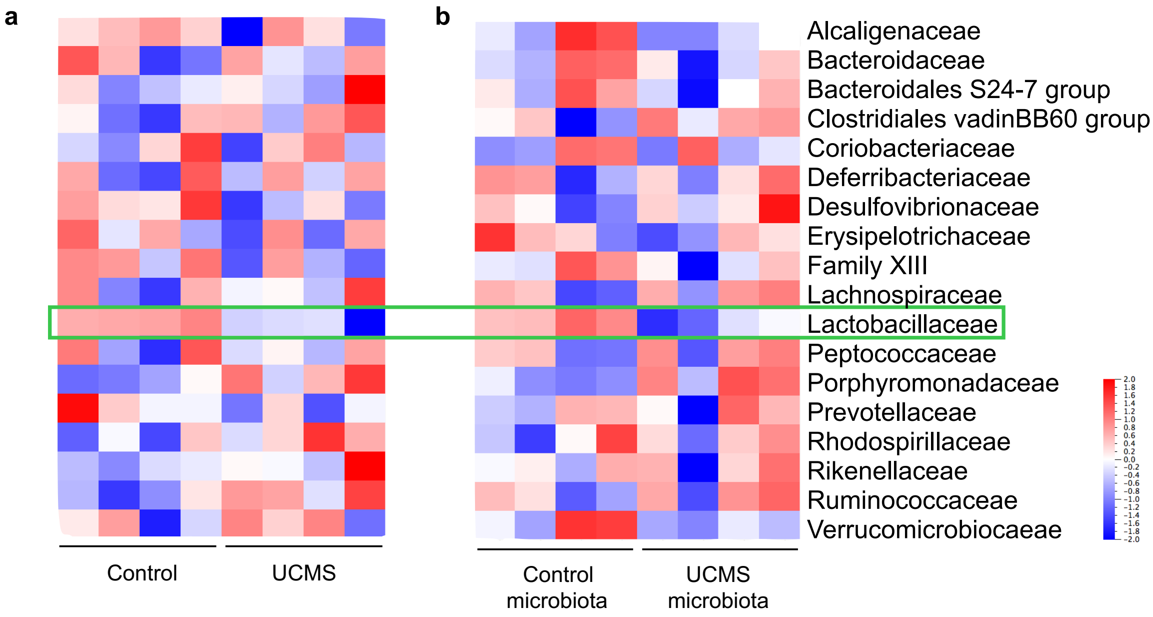


**Supplementary Figure 5 Unpredictable Chronic Mild Stress (UCMS) Alters Gut Microbiota and is Transferable to Recipient Mice.** (**a**,**b**) The 16S rDNA of the intestinal microbiota was sequenced and analyzed at the level of bacterial families in donor (*n* = 4/group) (**a**) and recipient mice (*n* = 4/group) (**b**).
